# Distinct roles of the two BRCA2 DNA binding domains in DNA damage repair and replication fork preservation

**DOI:** 10.1101/2024.09.24.614752

**Authors:** Francisco Neal, Wenjing Li, Mollie E. Uhrig, Neelam Sharma, Shahrez Syed, Sandeep Burma, Robert Hromas, Alexander Mazin, Eloise Dray, David Libich, Shaun Olsen, Elizabeth Wasmuth, Weixing Zhao, Claus S. Sørensen, Claudia Wiese, Youngho Kwon, Patrick Sung

**Affiliations:** Department of Biochemistry and Structural Biology, University of Texas Health at San Antonio, Texas 78229, USA; Greehey Children’s Cancer Research Institute, University of Texas Health at San Antonio, Texas 78229, USA; Department of Environmental and Radiological Health Sciences, Colorado State University, Fort Collins, Colorado, 80523, USA; Department of Neurosurgery, University of Texas Health at San Antonio, San Antonio, TX, 78229, USA; Department of Medicine, University of Texas Health at San Antonio, San Antonio, TX, 78229, USA; Biotech Research and Innovation Centre, Faculty of Health and Medical Sciences, University of Copenhagen, DK-2200 Copenhagen N, Denmark

## Abstract

Homologous recombination (HR) is a highly conserved tool for the removal of DNA double-strand breaks (DSBs) and the preservation of stalled and damaged DNA replication forks. Successful completion of HR requires the tumor suppressor BRCA2. Germline mutations in BRCA2 lead to familial breast, ovarian, and other cancers, underscoring the importance of this protein for maintaining genome stability. BRCA2 harbors two distinct DNA binding domains, one that possesses three oligonucleotide/oligosaccharide binding (OB) folds (known as the OB-DBD), and with the other residing in the C-terminal recombinase binding domain (termed the CTRB-DBD) encoded by the last gene exon. Here, we employ a combination of genetic, biochemical, and cellular approaches to delineate contributions of these two DNA binding domains toward HR and the maintenance of stressed DNA replication forks. We show that OB-DBD and CTRB-DBD confer ssDNA and dsDNA binding capabilities to BRCA2, respectively, and that BRCA2 variants mutated in either DNA binding domain are impaired in the ability to load the recombinase RAD51 onto ssDNA pre-occupied by RPA. While the CTRB-DBD mutant is modestly affected for HR, it exhibits a strong defect in the protection of stressed replication forks. In contrast, the OB-DBD is indispensable for both BRCA2 functions. Our study thus defines the unique contributions of the two BRCA2 DNA binding domains in genome maintenance.

## Introduction

Cells employ several mechanistically distinct tools for the elimination of DNA double-strand breaks (DSBs), lesions that can induce chromosome aberrations and rearrangements. One such tool is homologous recombination (HR), a high-accuracy process capable of faithfully restoring DNA integrity at DSB sites (San Filippo et al. 2008; Prakash et al. 2015; Scully et al. 2019). The HR machinery is also important for the removal of interstrand DNA crosslinks and the protection of stressed and damaged replication forks against nucleolytic attack (Cortez 2019; Tye et al. 2021). Given its broad involvement in genome maintenance, HR dysfunction can foster neoplastic transformation of cells and oncogenesis. A plethora of cancer-associated mutations have been identified in HR factors (Konstantinopoulos et al. 2015; Prakash et al. 2015; Zhao et al. 2019). In particular, somatic and germline mutations in the *BRCA1* and *BRCA2* genes, which encode proteins indispensable for HR and replication fork preservation, lead to genome destabilization and cancer in the breast, ovary, pancreas, and other organs (Konstantinopoulos et al. 2015; Prakash et al. 2015; Zhao et al. 2019).

Initiation of HR requires nucleolytic resection of the 5’ DNA strands at DSBs to generate 3’ ssDNA tails (Daley et al. 2015; Cejka and Symington 2021). These ssDNA tails then serve as the template for the assembly of a nucleoprotein filament of the recombinase RAD51, termed the presynaptic filament. The presynaptic filament mediates the search for an information donor sequence in a homologous chromatid and initiates invasion of the latter to form a DNA joint (Zhao et al. 2015; Zhao et al. 2019). Subsequent steps include repair DNA synthesis and resolution of DNA intermediates to yield repaired products (Zhao et al. 2019). Thus, presynaptic filament assembly represents a crucial step in the successful execution of HR. There is considerable evidence that formation of a RAD51-ssDNA nucleoprotein filament on a stressed or damaged replication fork that has undergone regression is necessary for fork protection against digestion by cellular nucleases (Cortez 2019; Zhao et al. 2019).

Presynaptic filament assembly is prone to interference by the single-strand DNA binding protein RPA (Bhat and Cortez 2018; Zhao et al. 2019). Owing to these constraints, cells have evolved mechanistically distinct protein factors, termed HR mediators, to facilitate RPA replacement by RAD51 on ssDNA (Filippo et al. 2006; Jensen et al. 2010; Zhao et al. 2015). BRCA2 (3,418 amino acid residues) functions in conjunction with its obligatory partner DSS1 (a 70-residue, highly acidic polypeptide) to mediate RPA-RAD51 exchange on ssDNA to facilitate presynaptic filament assembly (Zhao et al. 2015). BRCA2 provides the DNA binding and RAD51 interaction attributes of the BRCA2-DSS1 complex, while DSS1 serves as the RPA-recognition module (Zhao et al. 2015). Moreover, DSS1, through its association with a helical domain and OB fold 1 (HD-OB1), strongly attenuates the dsDNA binding activity of the OB-DBD (Zhao et al. 2015). Importantly, BRCA2 is also important for the protection of stressed replication forks against attack by nucleases such as MRE11 (Schlacher et al. 2011; Zhao et al. 2019; Kwon et al. 2023).

BRCA2 harbors two distinct classes of RAD51 interaction modules, namely, eight structurally related BRC repeats each capable of RAD51 association and the standalone C-terminal recombinase binding domain (CTRB), encoded by exon 27 (Lo et al. 2003; Carreira et al. 2009; Shivji et al. 2009; Chatterjee et al. 2016). Early studies identified a BRCA2 DNA binding domain that harbors three oligonucleotide/oligosaccharide binding (OB)-folds (Yang et al. 2002; Rajagopalan et al. 2010), which we refer to as the OB-DBD. Structural and biochemical evidence shows that OB folds 2 and 3 in the OB-DBD are directly involved in ssDNA binding (Yang et al. 2002; Rajagopalan et al. 2010). Importantly, our recent studies have uncovered a distinct DNA binding activity residing within the CTRB termed the CTRB-DBD (Kwon et al. 2023). Others have suggested the CTRB-DBD enables BRCA2 to diffuse along dsDNA in search of ssDNA to initiate RAD51 loading (Belan et al. 2023). Mutations that impair either the RAD51 interaction or DNA binding attribute of the CTRB exert a negative impact on HR mediator activity of BRCA2 *in vitro*, but only modest HR deficiency in cells and cellular resistance to DNA damaging agents (Kwon et al. 2023). However, simultaneous inactivation both CTRB attributes leads to more pronounced phenotypic consequences *in vitro* and in cells (Kwon et al. 2023). Separation-of-function CTRB mutations that specifically impair either RAD51 interaction (Schlacher et al. 2011; Kwon et al. 2023) or DNA binding (Kwon et al. 2023) affect the protection of replication forks. These findings help explain why deletion of BRCA2 gene exon 27, which encodes the CTRB, confers defects in HR, DNA damage repair, and cellular sensitivity to DNA damage (Sharan et al. 1997; Jasin 2002; McAllister et al. 2002; Donoho et al. 2003).

Here, we conduct biochemical and cellular analyses of OB-DBD and CTRB-DBD mutants to gain insights into the contributions of these two BRCA2 DNA binding domains to biological functions in HR, DNA damage repair, and replication fork preservation. Our studies reveal that while the OB-DBD is crucially important for optimal DNA damage repair and replication fork protection, the CTRB-DBD is more relevant for preserving the integrity of replication forks and fulfil a lesser role in DNA damage repair. Thus, our findings provide the first evidence for distinct roles of the two BRCA2 DNA binding domains in genome maintenance.

## Results and Discussion

### BRCA2-derived polypeptides for biochemical analyses

Previous studies have established that the HR mediator activity of BRCA2 can be interrogated using polypeptides that harbor key functional domains (**Figure 1A**). We have used BRC4-DBD (consisting of the BRC4 repeat fused to the OB-DBD) and miniBRCA2 (BRC4 fused to the OB-DBD and CTRB) (**Figure 1B**) in this study to define contributions of the OB-DBD and CTRB-DBD toward DNA binding affinities and specificities of BRCA2, as well as the HR mediator activity of BRCA2. These BRCA2 polypeptides were expressed in insect cells and purified to near homogeneity using procedures that we have devised (**Figure 1B**). We note that all BRCA2-derived polypeptides that harbor the OB-DBD were co-expressed with DSS1 and co-purified as stoichiometric complexes. This expression platform is equally suited to obtaining wild-type and mutant variants of the salient BRCA2 polypeptides (see **Methods**).

**Figure 1.**
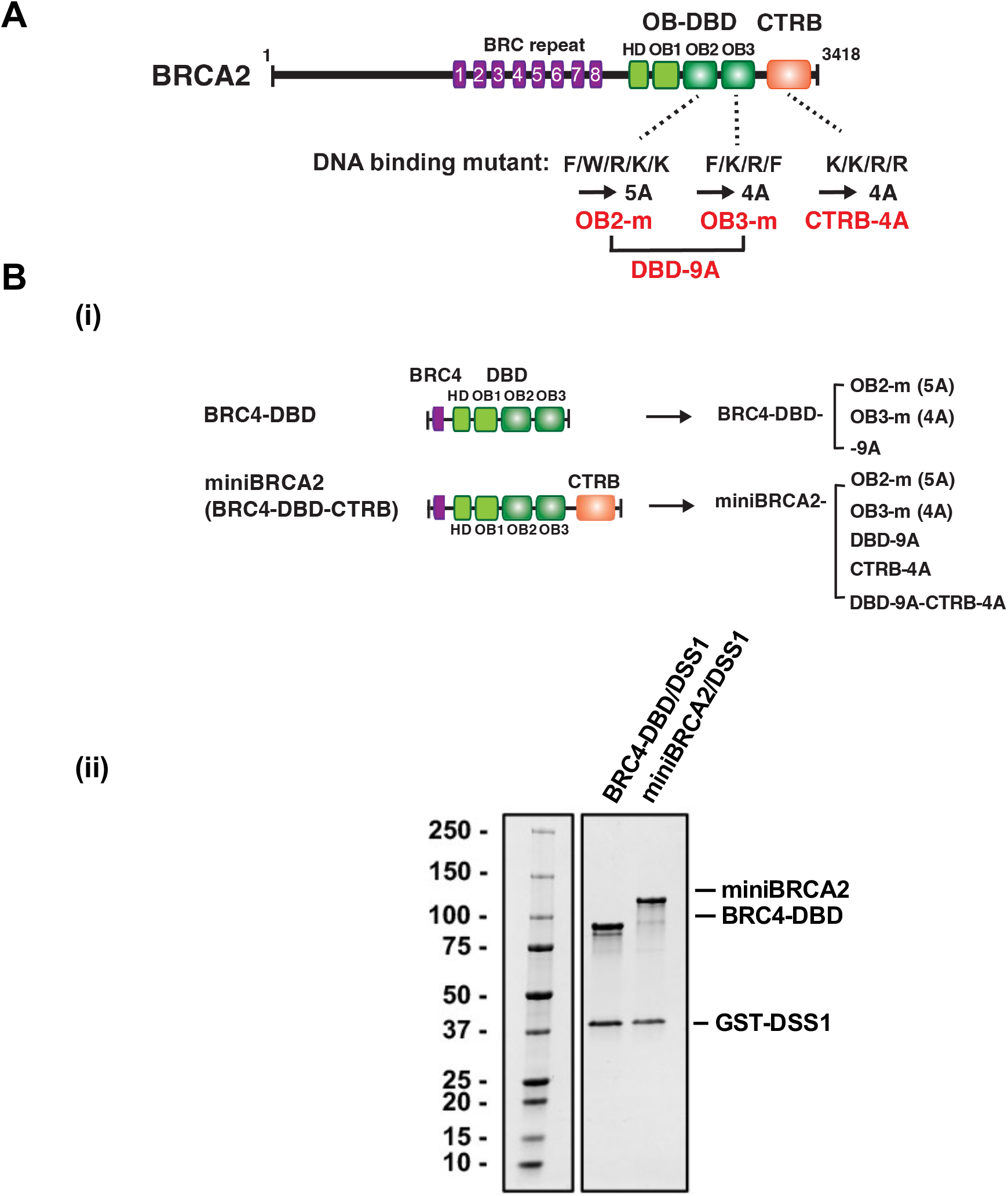
Mutagenesis of DBD and CTRB for DNA binding analysis. **(A)**Schematic of DNA binding domains in BRCA2 and mutations in DBD and CTRB. Residues in OB fold 2 (F2841, W2990. R2991, K3017, K3019), OB fold 3 (F3090, K3104, R3128, F3139), and CTRB (K3266, K3267, R3268, R3269) were mutated to Alanine. **(B)**Schematic of BRC4-DBD and BRC4-DBD-CTRB (miniBRCA2) and their mutant variants generated for biochemical studies **(i)**. SDS-PAGE showing purified wild-type BRC4-DBD/DSS1 and miniBRCA2/DSS1 **(ii)**. Purity analysis of mutants is presented in **Figure S1B**.

### Specificity of BRCA2 OB-DBD for ssDNA

As determined by X-ray crystallography, OB2 and OB3 of mouse Brca2 both directly contact ssDNA (Yang et al. 2002). In isolation, OB2 and OB3 of human BRCA2 bind ssDNA, and compound point mutations that impair the activity of these two OB folds have been described (Rajagopalan et al. 2010) (**Figure 1A, Figure S1A)**. Accordingly, we generated alanine substitution variants (OB2^5A^: F2841A, W2990A, R2991A, K3017A, and K3019A; OB3^4A^: F3090A, K3104A, R3128A, and F3139A; DBD^9A^: combining OB2^5A^ and OB3^4A^) within the context of the BRC4-DBD polypeptide derived from human BRCA2 (**Figure 1A, 1B**) and tested these mutant polypeptides alongside the wild-type counterpart for the binding of an 80-mer ssDNA substrate in an electrophoretic mobility shift assay (EMSA). The results showed that all three mutant BRC4-DBD polypeptides are significantly impaired for ssDNA binding activity (**Figure 2A**). Specifically, while the OB3^4A^ variant of BRC4-DBD retains residual ssDNA binding activity (about 40% at the highest concentration of 800 nM), the OB2^5A^ variant failed to bind the substrate even at the highest concentration (800 nM) tested (**Figure 2A**). Importantly, we found the DBD^9A^ mutant to be defective in ssDNA binding (**Figure 2A**). Our biochemical results thus demonstrate that both OB2 and OB3 contribute to ssDNA binding within the context of the OB-DBD.

**Figure 2.**
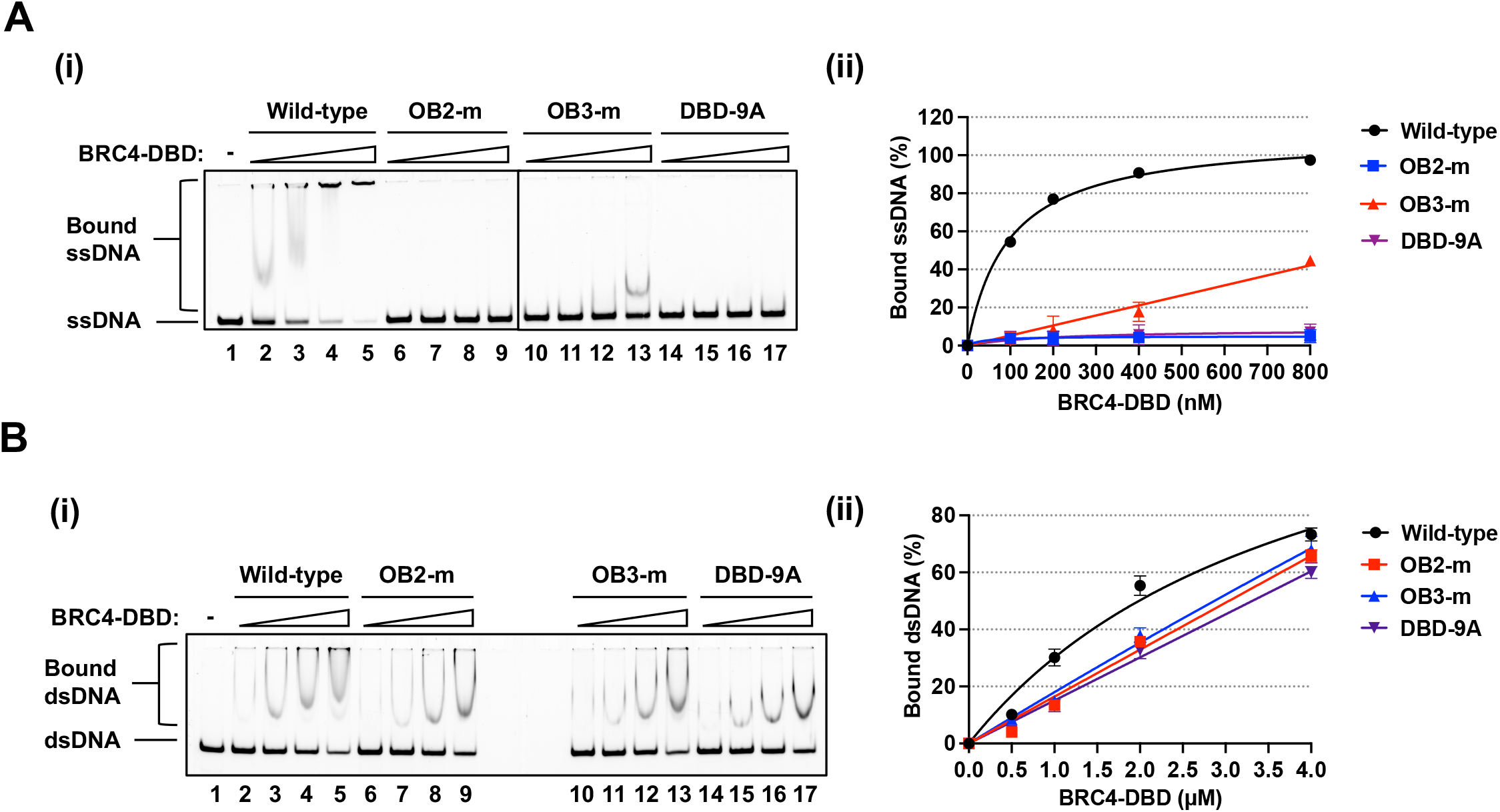
Role of OB2 and OB3 in ssDNA binding. **(A)**Results from EMSA testing BRC4-DBD WT, OB2-m, OB3-m, and DBD-9A. for ssDNA binding **(i)** and their quantification **(ii). (B)**Results from EMSA testing BRC4-DBD WT, OB2-m, OB3-m, and DBD-9A for dsDNA binding **(i)** and their quantification **(ii)**.

We next investigated whether BRC4-DBD in complex with DSS1 also has affinity for an 80-bp dsDNA. The results revealed that it has a much lower affinity for this substrate than for ssDNA (**Figure 2B**). While ∼90% ssDNA was bound by 400 nM of BRC4-DBD, ∼60% of the dsDNA was bound by as much as 2 µM of this polypeptide (**Figure 2B**). Even the DBD^9A^ mutation only engendered a modest defect in dsDNA binding (**Figure 2B**). These results indicate that the OB2 and OB3 DNA binding residues under study contribute more prominently to the recognition of ssDNA. We note that even though HD-OB1 within the OB-DBD has significant affinity for dsDNA, a recent study has shown that this activity is strongly attenuated by DSS1 and was therefore not detected in our analyses (Huang et al. 2024).

### Contributions of OB-DBD and CTRB-DBD to BRCA2 DNA binding affinity and specificity

Our prior biochemical analysis showed that the CTRB-DBD, in isolation, binds ssDNA and dsDNA with a comparable affinity and that a CTRB-DBD^4A^ (^3299^KKRR to A) mutation affects the binding to both substrates (Kwon et al. 2023). Here, we strived to determine the contributions of OB-DBD and CTRB-DBD to DNA binding affinity and specificity within the context of miniBRCA2. We tested miniBRCA2 variants that harbor salient OB and CTRB-DBD mutations to examine how they impact upon ssDNA and dsDNA engagement (**Figure 1B**). First, we determined that miniBRCA2 exhibits robust binding of ssDNA, with nearly all ssDNA substrate bound at 100 nM protein (**Figure 3A, Figure S3A**). All three OB fold variants show defective binding of ssDNA compared to wild-type miniBRCA2 (**Figure 3A, Figure S3A**), but they retain a higher degree of ssDNA binding compared to the corresponding BRC4-DBD mutants (**Figure 2A and 3A**). We surmise that the presence of intact CTRB-DBD within each OB fold mutant variant is responsible for the residual affinity for ssDNA in these miniBRCA2 species. The miniBRCA2 OB3^4A^ mutant shows a lesser ssDNA binding defect in comparison to the OB2^5A^ mutant (**Figure 3A(i), Figure S3A(i)**), which recapitulates the trend noted for the BRC4-DBD variants (**Figure 2A**). Taken together, these results support the premise that, within the context of miniBRCA2, the OB-DBD make a major contribution toward ssDNA engagement.

**Figure 3.**
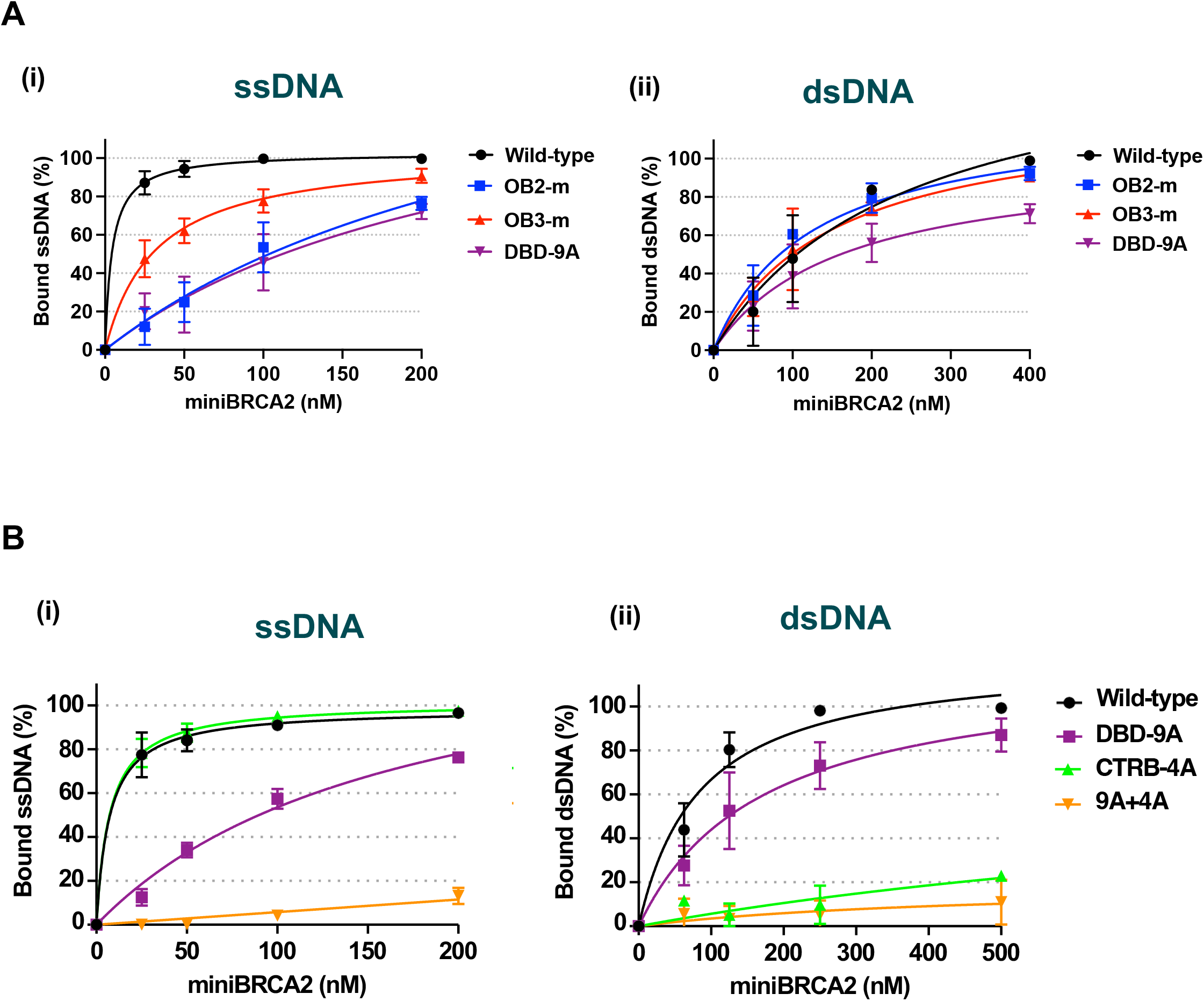
DNA binding by miniBRCA2 with OB fold and CTRB-DBD mutations. **(A)**Quantification of EMSA results for binding of ssDNA **(i)** and dsDNA **(ii)** by miniBRCA2 without or with OB2-m, OB3-m, or OB-DBD^9A^ mutation. Gel images are shown in **Figure S3A** and show a single, representative run for each indicated experimental comparison. **(B)**Quantification of EMSA results for binding of ssDNA **(i)** and dsDNA **(ii)** by miniBRCA2 bearing the OB-DBD^9A^, CTRB-DBD^4A^, or both mutations, alongside the wild-type protein. Gel images shown in **Figure S3B** illustrate a single, representative run for each indicated experimental comparison.

Our EMSA results show that miniBRCA2 binds dsDNA with avidity (**Figure 3A (ii), Figure S3A(ii)**). Given the OB-DBD has only weak affinity for dsDNA (**Figure 2B**), we posited that the CTRB-DBD represents the major dsDNA binding entity within miniBRCA2. Indeed, the miniBRCA2 OB fold 2 and 3 mutants exhibited comparable affinity for dsDNA relative to their wild-type counterpart, and even the OB-DBD^9A^ mutant only showed a modest deficiency in dsDNA binding (**Figure 3A(ii), Figure S3A(ii)**). These observations implicate the CTRB-DBD as the predominant dsDNA binding entity within miniBRCA2.

We examined the impact of the CTRB-DBD^4A^ mutation, alone or in combination with the OB-DBD^9A^ mutation, on ssDNA and dsDNA binding by miniBRCA2 (**Figure 1)**. This analysis confirmed that while the CTRB-DBD^4A^ mutation has no impact on ssDNA binding, it greatly affects the affinity for dsDNA (**Figure 3B, Figure S3B**). Notably, the OB-DBD^9A^/CTRB-DBD^4A^ double mutant is quite defective in binding either ssDNA or dsDNA (**Figure 3B, Figure S3B**). We also examined miniBRCA2 variants that harbor the OB-DBD^9A^ mutation, the CTRB-DBD^4A^ mutation, or both mutations for their affinity for a partially duplex substrate with a 3’ ssDNA overhang (**Figure S3C**). The results revealed moderate and little impairment of substrate binding by the OB-DBD^9A^ mutation and CTRB-DBD^4A^ mutation, respectively, while combining these mutations engendered a strong defect in substrate engagement (**Figure S3C**).

### Contributions of OB-DBD and CTRB-DBD to HR mediator activity

We showed previously that the CTRB-DBD^4A^ mutation exerts a negative impact on the HR mediator attribute of BRCA2-derived polypeptides that harbor the CTRB (Kwon et al. 2023). In this present study, we determined the effect of the OB-DBD mutations on HR mediator activity of the BRC4-DBD (**Figure S2B**) and miniBRCA2 (**Figure 4A**) polypeptides. In this analysis, RPA was added with RAD51 to a 167-mer ssDNA substrate to impede assembly of the presynaptic filament, followed by the addition of the indicated BRCA2 polypeptide and an incubation, before homologous duplex DNA was incorporated to initiate DNA strand exchange (see **Figure 4A(i)** for schematic). As expected, co-addition of RPA with RAD51 to ssDNA greatly suppressed the formation of the DNA strand exchange product (**Figure S2B(i; ii)**, lane 3). Importantly, while the addition of 500 nM of BRC4-DBD led to significant restoration of DNA strand exchange, the OB2^5A^, OB3^4A^, and OB-DBD^9A^ variants were all impaired for the HR mediator attribute, with the OB2^5A^ and OB-DBD^9A^ mutants exhibiting a more severe defect than the OB3^4A^ mutant in this regard (**Figure S2B(i; ii)**). The observed strand exchange defect is attributable to impaired DNA binding and not RAD51 interaction, as, by affinity pulldown, we showed that all three mutant BRC4-DBD polypeptides retain full ability to interact with RAD51 (**Figure S3D**).

**Figure 4.**
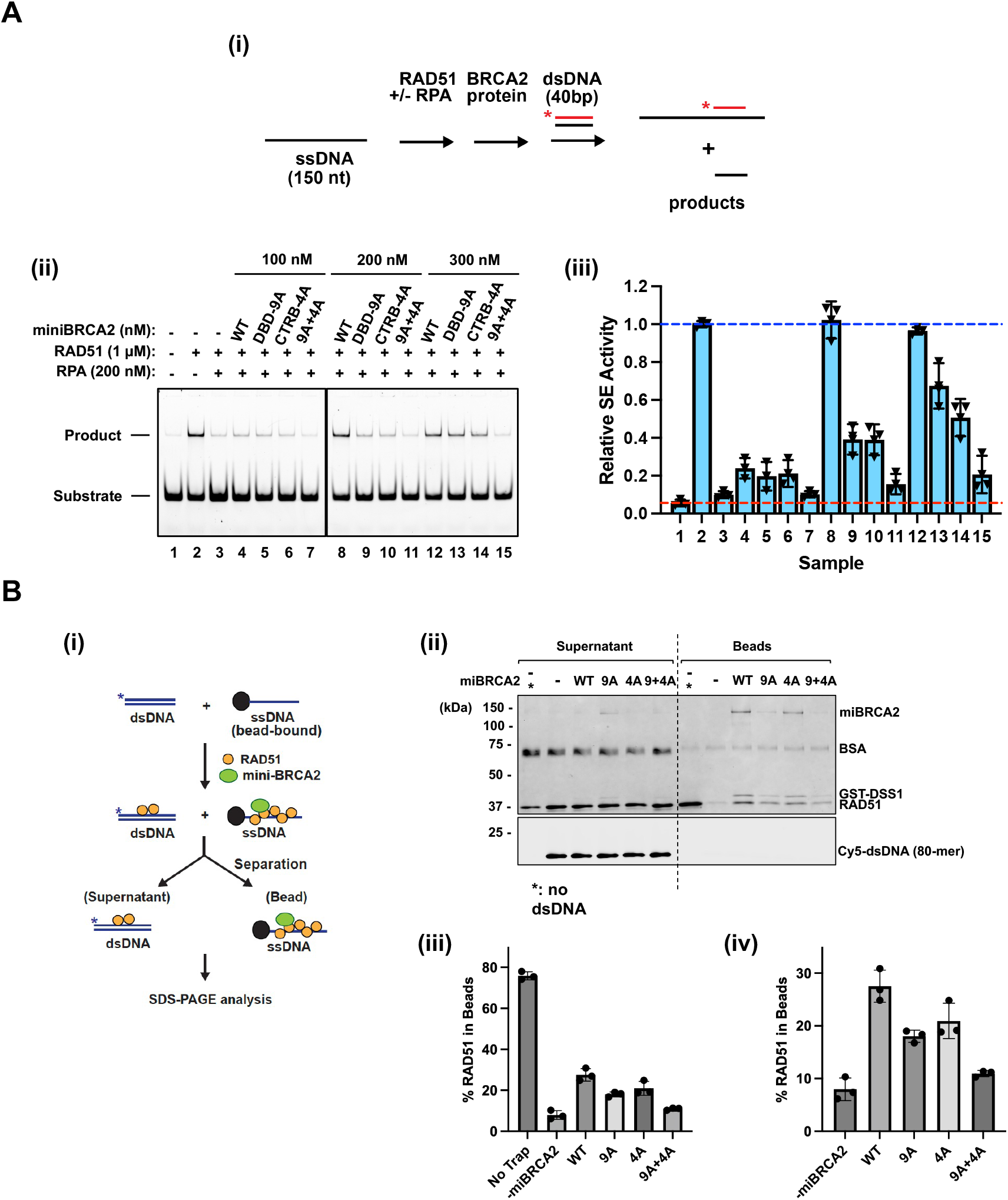
Roles of the OB-DBD and CTRB-DBD in ssDNA targeting of RAD51. **(A)**Strand exchange assay to test recombination mediator activity by miniBRCA2 and variants harboring the OB-DBD^9A^, CTRB-DBD^4A^, or the OB-DBD^9A^/CTRB-DBD^4A^ double mutation. The reaction schematic is shown in **(i)**, and results are presented in **(ii)**. Results were quantified **(iii). (B)**Schematic of RAD51 ssDNA targeting assay to test ability of miniBRCA2 (wild-type (WT) protein or the OB-DBD^9A^ (9A), CTRB-DBD^4A^ (4A), or OB-DBD^9A^/CTRB-DBD^4A^ (9+4A) mutant) to nucleate RAD51 onto ssDNA in the presence of a dsDNA trap **(i)**. SDS-PAGE to resolve proteins (Supernatant containing unbound proteins; Beads fraction with ssDNA-bound proteins) (**ii**). The dsDNA trap was revealed by Cy5 fluorescence **(ii)**. Results were quantified **(iii & iv)**. Panel **(iii)** shows percent of RAD51 trapped on the beads for all samples. Panel **(iv)** shows the same data, but with the Trap sample removed to better visualize the comparison of miniBRCA2 variants with each other and the miniBRCA2-negative sample.

Next, we tested the impact of the aforementioned OB fold mutations, alone or in combination with the CTRB-DBD^4A^ mutation, on the HR mediator attribute of miniBRCA2. Given that miniBRCA2 harbors both the OB-DBD and CTRB-DBD, it is significantly more efficacious than BRC4-DBD in restoring DNA strand exchange upon co-incubation of the ssDNA template with RPA and RAD51. Specifically, 200 nM of miniBRCA2 (**Figure 4A(ii; iii)**), in contrast to 500 nM of BRC4-DBD (**Figure S2B**), is needed for significant restoration of DNA strand exchange. Importantly, at 200 nM concentration of miniBRCA2, the OB-DBD^9A^ mutation imparts as strong a deficiency in HR mediator activity as the CTRB-DBD^4A^ mutation and combining these two mutations further compromises mediator activity (**Figure 4A(ii; iii), Figure S2B)**. The same general trend in HR mediator efficacy was seen when the miniBRCA2 variants were tested at 300 nM concentration (**Figure 4A(ii; iii))**.

### Impact of OB-DBD and CTRB-DBD mutations in ssDNA targeting of RAD51

We previously showed that the CTRB facilitates nucleation of RAD51 onto ssDNA, a critical step in generating the presynaptic filament (Kwon et al. 2023). Here, we sought to elucidate what contributions the two BRCA2 DNA binding domains make in RAD51 targeting to ssDNA for subsequent nucleoprotein filament assembly. We utilize an *in vitro* experimental protocol (**Figure 4B(i)**) to interrogate how BRCA2 polypeptides nucleate RAD51 onto ssDNA in the presence of a dsDNA trap. We employed this analytical tool to elucidate how loss of DNA binding by the OB-DBD or CTRB-DBD would impair nucleation of RAD51 onto ssDNA (**Figure 4B (i)**). We found that miniBRCA2 variants that harbor either the OB-DBD^9A^ or CTRB-DBD^4A^ mutation are only moderately impaired in RAD51 ssDNA-targeting (**Figure 4B (ii)**), but the mutant polypeptide that harbors both mutations is quite defective in this regard (**Figure 4B(ii-iv)**).

### Role of the OB-DBD and CTRB-DBD in DNA damage repair and HR

For testing the impact of the OB-DBD mutations on the biological functions of BRCA2, we generated DLD1 cells stably expressing the salient mutants in full-length BRCA2 protein. We transfected DLD1 cells (BRCA2^-/-^) with the salient expression vectors and then isolated stable clones expressing equivalent levels of wild-type, OB-DBD^9A^ mutant, and CTRB-DBD^4A^ mutant versions of BRCA2 (**Figure 5A and Figure S4A**). We note that all three ectopically expressed BRCA2 species localize to the nucleus properly (**Figure S4A**). Notably, cells expressing the OB-DBD^9A^/CTRB-DBD^4A^ double mutant show severe growth defects and cell morphology changes (**Figure S4B)**. These poor growth characteristics of the double mutant cells rendered them unsuitable for cell biological studies. This observation provides evidence that simultaneous inactivation of the two DNA binding domains in BRCA2 is likely incompatible with cell viability. Thus, we focused our cellular studies primarily on the OB-DBD^9A^ and CTRB-DBD^4A^ single mutants.

**Figure 5.**
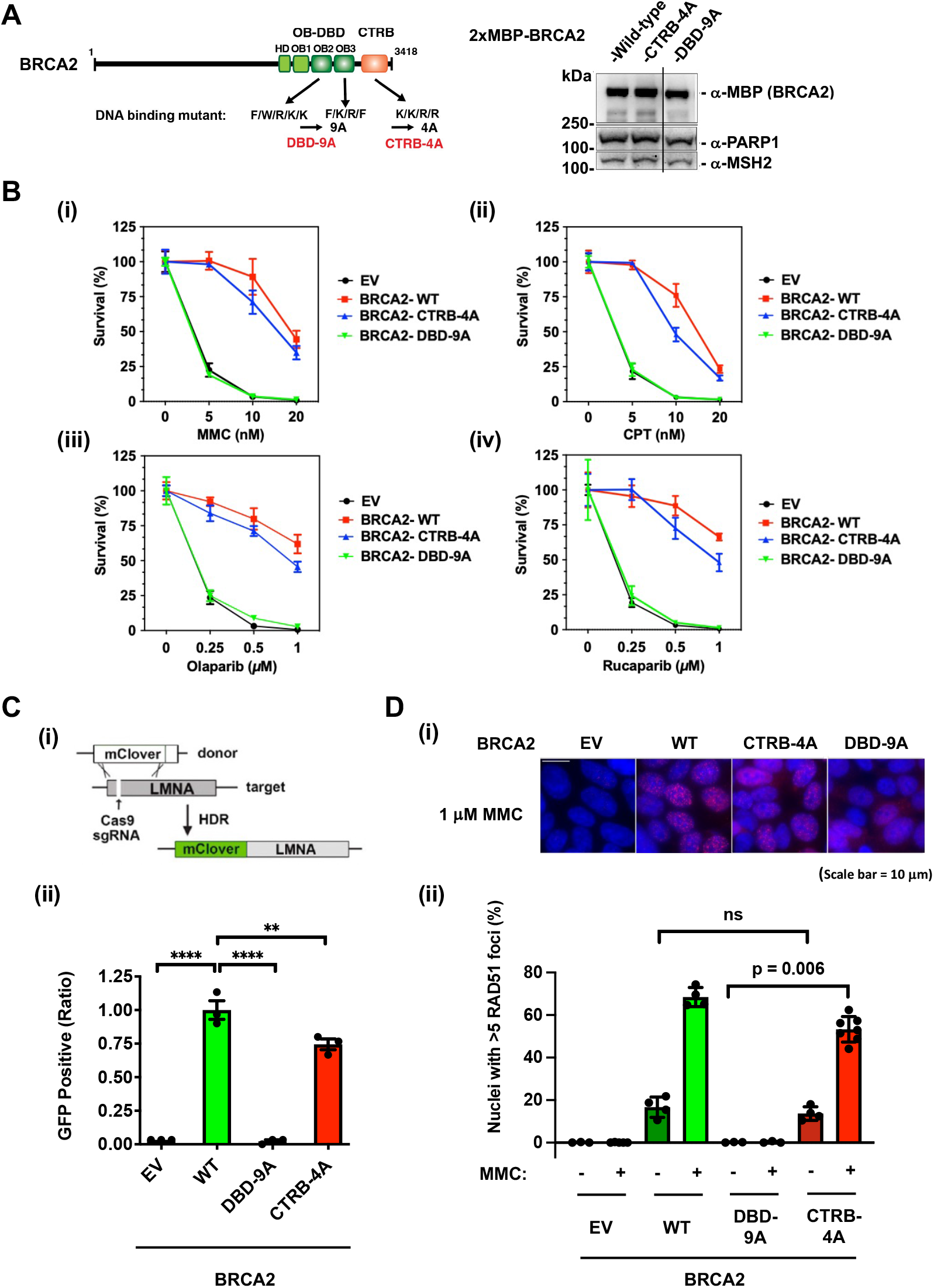
Mutation of the DBD compromises cellular HR function. **(A)**Schematic of the OB-DBD^9A^ and CTRB-DBD^4A^ mutant proteins tested in cell-based analyses. Western blot of nuclear fractions of DLD1 cells expressing the BRCA2 proteins. All three BRCA2 species are present in the nuclear fraction (see **Figure S4A**). **(B)**Clonogenic survival of DLD1 cell lines upon treatment with mitomycin C (MMC) **(i)**, camptothecin (CPT) **(ii)**, and the PARP inhibitors olaparib **(iii)** and rucaparib (**iv**). **(C)**Schematic of CRISPR/Cas9 gene integration assay used to test HR proficiency of DLD1 cell lines **(i)**. Results are shown in **(ii). (D)**RAD51 focus formation was accessed with or without exposure to MMC. Shown are representative micrographs of RAD51 foci (**i**) and quantified results (**ii**). Representative micrographs of untreated cells are presented in **Figure S4C**.

With the requisite cell lines in hand, we first sought to determine the impact of DBD and CTRB mutations on the sensitivity of cells to several DNA-damaging agents, namely camptothecin (CPT), mitomycin C (MMC), and the PARP inhibitors olaparib and rucaparib (**Figure 5B**). This analysis revealed that the OB-DBD^9A^ mutation engenders a high degree of sensitivity to all four DNA-damaging agents, being equivalent to the hypersensitivity observed for control DLD1 cells (BRCA2^-/-^) transfected with the empty vector (**Figure 5B**). In comparison, and in concordance with what we have reported recently (Kwon et al. 2023), the CTRB-DBD^4A^ mutation engenders only mild DNA damage sensitivity to the same panel of DNA damaging agents (**Figure 5B**). These results suggest that DNA binding by the OB-DBD plays a crucial role in DNA damage repair and ultimately supports cell viability upon the occurrence of a variety of genomic insults.

Next, we used a CRISPR/Cas9-based gene targeting assay (Pinder et al. 2015) to interrogate how the OB-DBD and CTRB-DBD mutations affect HR efficiency (**Figure 5C**). In agreement with our previous studies (Kwon et al. 2023), we found that DLD1 cells expressing the CTRB-DBD^4A^ mutant exhibit a modest decrease in HR efficiency (**Figure 5C**). Importantly, DLD1 cells expressing the OB-DBD^9A^ mutant are markedly impaired in HR (**Figure 5C**), which parallels the striking sensitivity to DNA-damaging agents observed for these mutant cells. We also determined the ability of DLD1 cells of different genetic lineages to assemble nuclear RAD51 foci upon MMC treatment (**Figure 5D, Figure S4C**). In concordance with results from other cellular analyses (**Figure 5B** and **Figure 5C**), the OB-DBD^9A^ mutation eliminates formation of RAD51 foci in response to treatment with mitomycin C (MMC), while CTRB^4A^ mutation mildly affects focus formation in the treated cells (**Figure 5D, Figure S4C**). Our results thus reveal a prominent role for ssDNA binding via OB-DBD to drive HR-mediated repair of DNA damage, while DNA binding by the CTRB-DBD makes a lesser contribution in this regard.

### Involvement of OB-DBD and CTRB-DBD in replication fork preservation

BRCA2 fulfills a prominent role in the protection of stressed replication forks against nucleolytic attrition, predominantly by the MRE11 nuclease (Kim et al. 2014; Mijic et al. 2017; Halder et al. 2022). Here, we used the DNA fiber assay to determine the role of the two BRCA2 DNA binding domains in the protection of replication forks upon treatment of cells with hydroxyurea (HU), a potent replication stressor. Cells were sequentially pulsed with CIdU and IdU before being exposed to 2 mM HU for 5 h, and IdU/CldU tract ratios were determined to assess the extent of fork degradation (**Figure 6A (i), Figure S4D**). As shown in **Figure 6A (ii)**, parental DLD1 cells (BRCA2^-/-^) or DLD1 cells ectopically expressing wild-type BRCA2, the OB-DBD^9A^ mutant, or CTRB-DBD^4A^ show no significant difference in IdU/CldU tract ratios in the absence of HU. Importantly, cells expressing either the OB-DBD^9A^ or CTRB^4A^ mutant are highly prone to replication fork attrition upon exposure to 2 mM HU (**Figure 6A(ii**)). We also verified that replication fork attrition in either OB-DBD^9A^ or CTRB^4A^ mutant cells could be alleviated to a large degree by treatment with the MRE11 nuclease inhibitor Mirin (**Figure 6A(ii**)). Collectively, these results support the notion that both DNA binding domains of BRCA2 are indispensable for the protection of stressed replication forks against nucleolytic attack.

**Figure 6.**
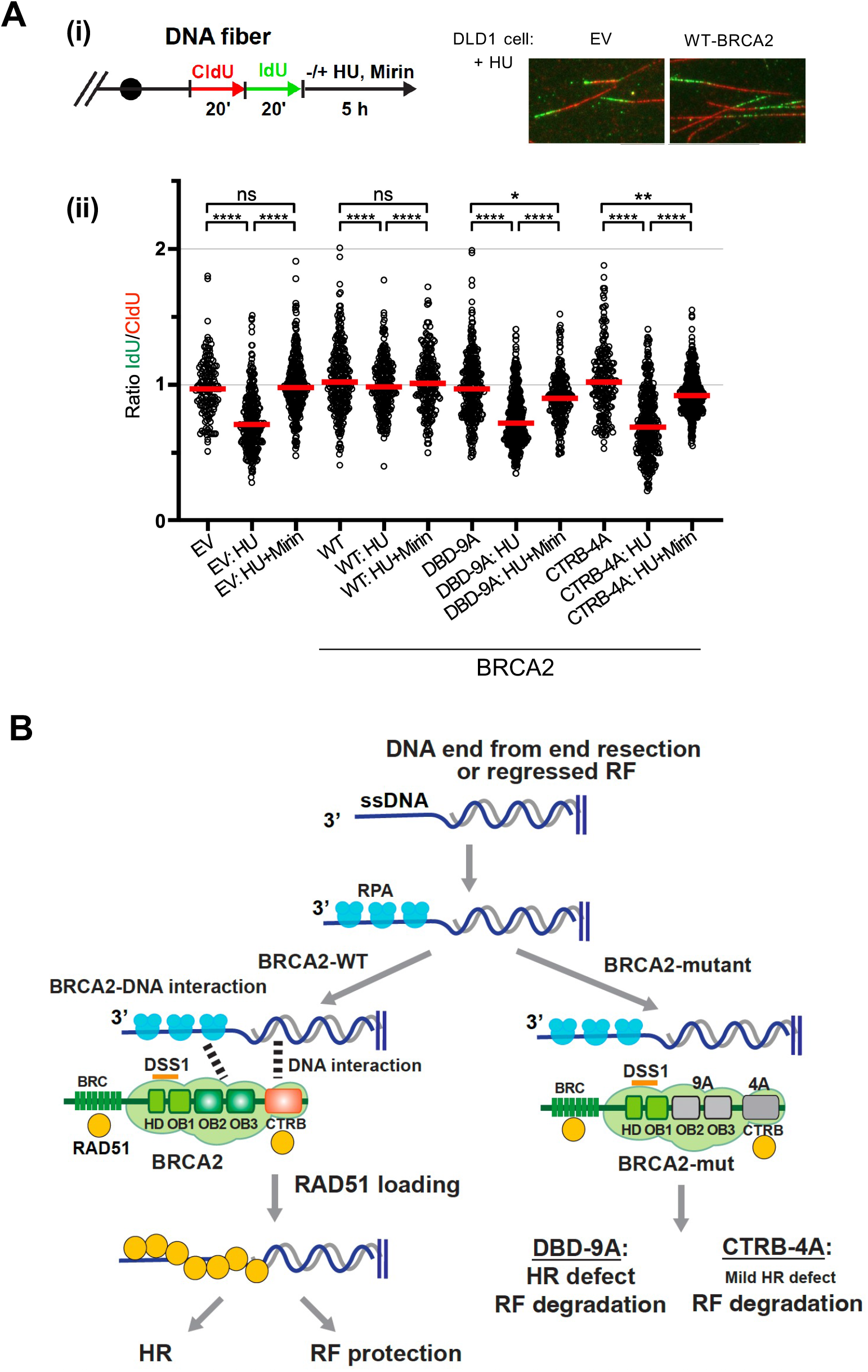
Impact of OB-DBD and CTRB-DBD mutations on replication fork preservation. **(A)**Overview of DNA fiber assay to test replication fork protection in DLD1 cell lines and representative micrographs of CldU/IdU-labeled DNA fibers **(i)**. Additional micrographs are presented in **Figure S4D**. Dot plots of IdU/CldU tract length ratios in DLD1 cells and derivatives left untreated, treated with 2 mM hydroxyurea (HU), or 2 mM HU and 50 μM Mirin **(ii). (B)**Model showing contributions of the BRCA2 DNA interaction via DBD and CTRB toward DNA repair by HR or replication fork protection, and the outcome of the mutations disrupting the interactions.

### Synopsis and mechanistic model for BRCA2 functions

The role of BRCA2 in DSB repair and replication fork preservation is well-established (Konstantinopoulos et al. 2015, Prakash et al. 2015, Zhao et al. 2019). However, how this tumor suppressor protein utilizes its OB-DBD and CTRB-DBD domains (Yang et al. 2002, Zhao et al. 2019, Kwon et al. 2023) to orchestrate repair and replication fork protection remains to be defined. Here, we have addressed not only the contributions of these two BRCA2 DNA binding domains in ssDNA and dsDNA engagement, but also their functional significance in RAD51 presynaptic filament assembly and in DNA damage repair, HR, and the protection of stressed replication forks against nucleolytic attrition within the cellular setting. Specifically, by constructing multiple OB fold mutants within the context of functional BRCA2 polypeptides and testing them in a variety of reconstituted biochemical systems and in cell-based assays, we have provided evidence that the OB-DBD is primarily involved in ssDNA engagement and plays a more prominent role than the CTRB-DBD in RAD51 presynaptic filament assembly. Accordingly, mutations in the OB-DBD that ablate ssDNA binding compromises the DNA damage repair functions of BRCA2 in cells. We also provide evidence that inactivation of the CTRB-DBD DNA binding attribute leads to only a modest reduction in RAD51 presynaptic filament assembly *in vitro* and in HR and DNA damage repair efficiency in cells. However, the CTRB-DBD mutation under study engenders a replication fork protection phenotype just as severe as the compound OB-DBD mutation that simultaneously ablates OB2 and OB3 function. Altogether, our results show that the OB-DBD and CTRB-DBD are not redundant entities, but, rather, serve distinct biological roles, with the former being indispensable for both HR execution and replication fork protection, and the latter as being particularly germane for replication fork preservation. A model depicting the distinctive roles of the OB-DBD and CTRB in the functional engagement of HR and replication fork intermediates is presented in **Figure 6B**.

### Methods

#### Plasmids

We introduced cDNA encoding BRC4-DBD or miniBRCA2 (**Figure 1, Table 1**) into the MacroBac vector p438a (Gradia et al. 2017). BRC4-DBD and miniBRCA2 were co-expressed with GST-tagged DSS1 in Hi5 insect cells (*Tni* cells; Expression Systems). For cellular studies, full-length BRCA2, with an N-terminal 2xMBP tag and a C-terminal His6-Flag tag, was introduced into phCMV1 (**Table 1**). Site-directed mutagenesis (Gene Universal and Twist Bioscience) was used to generate mutations in the DBD, CTRB, or both (**Table 2**) into the indicated BRC4-DBD, miniBRCA2, and full-length BRCA2 constructs.

#### Mammalian Cell Culture

DLD1 cells (BRCA2^-/-^, Horizon Discovery) were maintained in RPMI 1640 media (Gibco, Catalog #11875-093). To express full-length BRCA2, DLD1 cells were transfected with 2 µg of pCMV-2xMBP-BRCA2-His6-Flag (wild-type BRCA2 or the indicated mutant) using lipofectamine 2000 (Invitrogen) according to the manufacturer’s directions. Single colonies stably expressing the indicated full-length BRCA2 constructs were selected and maintained using G418 (0.5 mg/ml). DLD1 cell lines were grown at 37 °C and 5% CO_2_.

#### Expression and purification of recombinant proteins

Lysate preparation and protein purification steps were carried out between 0-4 °C. Purified proteins were concentrated to a small volume in an Amicon ultracentrifugation device (Millipore), snap frozen in 2-4 µl aliquots, and stored at -80 °C. Once thawed, proteins were kept on ice and unused fractions were discarded after 48 h.

#### BRC4-DBD and miniBRCA2 complexed with DSS1

Recombinant bacmid DNA encoding BRC4-DBD, miniBRCA2, or DSS1 was generated from DH10Bac cells (Thermo Fisher Scientific) using p438a-BRC4-DBD or miniBRCA2, or pDEST20-DSS1. Baculoviruses were amplified in SF9 cells (Thermo Fisher Scientific). Hi5 insect cells were co-transfected with baculovirus encoding BRC4-DBD or miniBRCA2 and baculovirus encoding DSS1 at 27 °C for 48 h. Cells were harvested by centrifugation, and pellets were stored at -80 °C. Cell lysate was prepared by using a Dounce homogenizer to resuspend the Tni cell pellets in buffer T (25 mM Tris-HCl, pH 7.5, 10% glycerol, 0.5 mM EDTA, 1 mM DTT, 0.01% Igepal CA–630) containing 300 mM KCl, 1 mM benzamidine, protease inhibitors (2 μg/ml each of aprotinin, chymostatin, leupeptin, and pepstatin A). Lysate was clarified by centrifugation at 40,000 rpm x 45 min x 4 °C. The supernatant was incubated with 2 ml anti-Flag M2 affinity gel (Sigma-Aldrich) in lysis buffer for 2 h at 4 °C on a rocking shaker. Resin was washed with 3 ml each of T buffer containing 300 mM KCl, 1000 mM KCl with 1 mM ATP and 8 mM MgCl_2_, and finally 100 mM KCl. Protein was eluted from resin using buffer T containing 100 mM KCl and 0.5 mg/ml Flag peptide. Fractions of eluted protein were pooled and mixed with 2 ml of Glutathione Sepharose 4 Fast Flow (Cytiva) for 1.5 h on a rocking shaker. Glutathione resin was washed in buffer T containing 100 mM KCl, 1000 mM KCl with 1 mM ATP and 8 mM MgCl_2_, and finally 100 mM KCl. Protein was eluted off glutathione resin using buffer T containing 100 mM KCl and 50 mM glutathione, pH 8.0. Eluted protein fractions were pooled, diluted 2-fold in buffer T, and loaded into a 1 ml Mono-Q column (GE Healthcare). Bound protein was eluted using a salt gradient ranging from 50 mM KCl to 350 mM KCl. Peak protein fractions containing BRC4-DBD/DSS1 or miniBRCA2/DSS1 were pooled and concentrated using a 30K Amicon Ultra centrifugal device (Millipore). Concentrated protein was aliquoted, snap frozen in liquid nitrogen, and stored at -80 °C. This purification scheme was used to purify wild-type and all mutant variants of BRC4-DBD/DSS1 and miniBRCA2/DSS1. Throughout each purification, BRC4-DBD/DSS1 and miniBRCA2/DSS1 were verified by western blot with anti-Flag HRP antibody (Invitrogen) for BRCA2 peptides and anti-GST HRP conjugate (GE Healthcare) for DSS1.

#### RAD51

We expressed His 6-Smt3-RAD51 using *E coli* Rosetta cells (Sigma-Aldrich) (Kaminski et al. 2022). Bacterial cells were cultured in 2X luria broth (supplemented with 100 µg/ml ampicillin) and induced by incubation with 0.5 mM IPTG for 16 h at 37 °C. Bacterial cells were harvested by centrifugation (5000 rpm x 10 min), and pellets were stored at -80 °C. For cell lysis, pellets were resuspended in buffer T containing 600 mM KCl, 6 U/µl benzonase, 1 mM MgCl_2_, and protease inhibitors (1 mM benzamidine and 2 μg/ml each of aprotinin, chymostatin, leupeptin, and pepstatin A) and then subject to sonication. Lysate was clarified by centrifugation (45,000 rpm x 45 min). Supernatant from centrifugation was combined with (NH4)_2_SO_4_ (0.277 g per ml lysate) to precipitate protein. The precipitate was harvested by centrifugation 18,000 x *g* for 30 min. The pellet was dissolved in buffer T (containing 300 mM KCl) and incubated with 2 ml Ni-NTA Superflow resin (QIAGEN) for 2 h on a rocking shaker. The resin was washed with buffer T containing 500 mM KCl, 300 mM KCl, and finally 150 mM KCl. To remove the His6-Smt3 tag, RAD51 bound to resin was digested with Ulp1 (5 µg) in buffer T (150 mM KCl) for 16 h at 4 °C. RAD51 released from the resin was collected in buffer T containing 150 mM KCl. After adjusting the salt concentration to 150 mM KCl, the protein pool was fractionated in a 1 ml HiTrap Q HP column and eluted over a gradient of 100 to 500 mM KCl. Peak fractions containing protein were pooled, concentrated, and further fractionated in a 1 ml Heparin HP column, which was eluted with a salt gradient of 100 to 500 mM KCl. Peak fractions containing protein were pooled, concentrated, and stored.

#### RPA

We expressed and purified RPA using our published protocol (Sigurdsson et al. 2001).

#### DNA substrates

HPLC-purified DNA oligonucleotides (**Table 3**) were purchased from IDT. Blunt-ended dsDNA substrate and partial dsDNA substrate with a 3’ ssDNA overhang were generated by annealing equimolar amounts of the indicated oligonucleotides as described (Kwon et al. 2011).

#### Electrophoretic mobility shift assay

BRCA2-derived polypeptides (BRC4-DBD and miniBRCA2) were incubated with 5 nM of the indicated Cy5-labeled DNA substrate (ssDNA, dsDNA, partial duplex DNA with a 3’-overhang) (**Table 3**) in reaction buffer (35 mM Tris, pH 7.5, 1 mM DTT, 50 mM KCl, 1 mM MgCl_2_, and 100 μg/ml BSA) for 10 min at 25 °C. Unbound DNA and nucleoprotein complexes were resolved by electrophoresis on 8% polyacrylamide gels in TBE buffer (50 mM Tris-boric acid, pH 8.4, 0.5 mM EDTA) for 1 h at 40 mA. DNA species were visualized using the ChemiDoc imaging system (Bio-Rad). The proportion of bound versus unbound DNA was quantified using the Image Lab software (Bio-Rad). Mean bound DNA and standard deviation were calculated across 3 independent studies using Microsoft Excel and GraphPad Prism 9. Data were plotted as means on an X-Y scatterplot with a best fit curve, with error bars representing SD for each experimental sample.

#### DNA strand exchange assay

Unless stated otherwise, reaction steps were carried out at 37 °C. To test for recombination mediator activity of BRCA2 polypeptides, strand exchange reactions were assembled (12.5 µl final volume) using 20 nM ssDNA (oligo 863 in **Table 3**) in buffer B (35 mM Tris, pH 7.5, 1 mM DTT, 50 mM KCl, 1 mM ATP, 1 mM MgCl_2_, and 100 μg/ml BSA). The ssDNA was incubated with 1 µM RAD51 and 200 nM RPA for 10 min before BRC4-DBD/DSS1 or miniBRCA2/DSS1 was added to the indicated concentrations. After a 5 min incubation, strand exchange was initiated by adding 20 nM of duplex DNA (5’ Cy5-labeled oligo 1056 with oligo 1057, **Table 3**) and 4 mM spermidine hydrochloride. Reactions were incubated for 30 min and mixed with 1 µl each of 1% SDS and proteinase K (10 mg/ml). Following a 5 min incubation, reaction mixtures were resolved in 8% polyacrylamide gels in TBE buffer (50 mM Tris-borate, pH 8.4, 0.5 mM EDTA). Cy5-labeled substrate and strand exchange product were visualized in the ChemiDoc imaging system (Bio-Rad). Proportion of product DNA formed was quantified and compared relative to the RAD51-only sample (lane 2). Mean relative strand exchange (SE) activity compared to RAD51-only activity was quantified across 3 separate studies using GraphPad Prism 9 and Microsoft Excel. Mean SE activity was plotted with individual data points and error bars (SD) shown for each experimental sample.

#### RAD51 ssDNA targeting assay

To assemble the reactions (final volume of 20 µl), 2.5 µM RAD51 was incubated with 0.25 µM miniBRCA2 in buffer C (35 mM Tris-HCl, pH 7.5, 1 mM MgCl_2_, 1 mM ATP, 50 mM KCl, and 50 ng/μL BSA) for 10 min at 4 °C. Then, 200 nM of duplex DNA (5’ Cy5-labeled oligo 2040 with oligo 2041, **Table 3**) and 50 nM of ssDNA (biotin-dT80)-immobilized on 4 µl Dynabeads M270 Streptavadin (Invitrogen) was added to each reaction followed by a 10 min incubation at 25 °C. Dynabeads were captured using a Magnetic Particle Separator (Roche Applied Science), which were washed twice in buffer containing 35 mM Tris-HCl, pH 7.5, 1 mM MgCl_2_, 0.1 mM ATP, and 50 mM KCl. Bound protein was eluted at 37 °C for 3 min with 10 µl of SDS-PAGE loading buffer. The supernatant and eluate fractions were analyzed by SDS-PAGE with Coomassie blue staining. Following electrophoresis, Cy5-labeled dsDNA was visualized in the ChemiDoc imaging system. Mean proportion of RAD51 collected in eluate fractions was calculated for 3 separate experiments. GraphPad Prism 9 was used to tabulate SD and construct graphs depicting mean % RAD51 in eluate samples, with SD represented in accompanying error bars.

#### Affinity pulldown assay

The indicated BRCA2 polypeptide (2 µg) was incubated with RAD51 (6 µg) in a 25 µl reaction assembled in buffer T (150 mM KCl) and 6 U/µl benzonase for 30 min on ice. Following incubation, reactions were mixed with anti-Flag M2 affinity gel and incubated on a rocking shaker for 1 h at 4 °C. Resin was washed three times with 100 µl buffer T (150 mM KCl) and then incubated with 10 µl of SDS-PAGE loading buffer at 37 °C for 5 min to elute proteins. The supernatant and eluate fractions were analyzed by SDS-PAGE and Coomassie blue staining.

#### Cell fractionation

The Rapid, Efficient And Practical (REAP) method for the preparation of cytoplasmic and nuclear extracts was followed (Suzuki et al. 2010). Briefly, DLD1 cells on 10-cm dishes were washed with ice-cold phosphate buffer saline (PBS) pH 7.4, collected by centrifugation, resuspended in 900 µl of ice-cold PBS with 0.05% NP40 and protease inhibitors, and triturated 5 times using a p1000 micropipette. The lysed cell suspension was centrifuged to separate the supernatant (representing the cytoplasmic fraction), and the pelleted nuclei were washed once with PBS containing 0.05% NP40 and lysed with NETN buffer (20 mM Tris-HCl pH 8, 420 mM NaCl, 1 mM EDTA, 0.5% Igepal CA630, 1 mM DTT) with protease inhibitors to yield the nuclear fraction. The cytoplasmic and nuclear fractions, 20 or 40 µg each, were analyzed by immunoblotting.

#### Clonogenic survival assay

DLD1 cells expressing full-length BRCA2 (wild-type or the indicated DNA binding mutant), or empty vector (phCMV1-2xMBP), were seeded in 6 well plates at 300 cells per well and incubated for 24 h at 37 °C. The next day, cells were treated for 12 days with the indicated concentration of camptothecin (CPT), mitomycin C (MMC), olaparib, or rucaparib. Following drug treatment, cells were fixed in 100% methanol, then stained in 0.5% crystal violet in 20% methanol to visualize DLD1 colonies. To determine clonogenic survival for each drug treatment, the number of surviving colonies in experimental plates was compared to the number of surviving colonies in untreated control plates.

#### CRISPR-Cas9 gene targeting assay

DLD1 cells expressing 2xMBP-BRCA2-His6-Flag (wild-type or DNA binding mutant) or empty vector were seeded in 6-well plates at 4 × 10^5^ cells per well. Cells were incubated 37 °C for 24 h and then were transfected with 4 µl lipofectamine 2000 (Invitrogen), and a 1.5:1 or 2:1 µg ratio of sgRNA plasmid px330-LMNA to pCR2.1-CloverLamin donor template (Pinder et al. 2015). After a 72 h incubation with gene transfer plasmids, cells were collected in 300 µ PBS containing 5% FBS for flow cytometry analysis. Cells positive for GFP were detected by flow cytometry, and the mean proportion of GFP+ cells in each sample was quantified across 3 independent experiments. Data are presented as the mean percent of GFP+ cells normalized to % of GFP+ of BRCA2 WT, with accompanying error bars signifying the standard deviation.

#### RAD51 foci

Eight-well Permanox chamber slides (LabTek) were seeded with 40,000 cells per chamber 48 h before treatment with 1 μM MMC for 16 h. Following treatment, cells were washed twice with PBS, fixed in 1% paraformaldehyde/2% sucrose in PBS for 15 min at room temperature, washed twice with PBS and permeabilized with methanol for 30 min at –20 °C, as described (Jimenez-Sainz et al. 2022). Cells were washed twice in PBS before incubation in 0.5% TritonX-100 /PBS for 10 min. Samples were blocked in 5% BSA/ PBS for 30 min at 25 °C before incubation with primary antibody (⍰-RAD51 from BioAcademia 70-001, 1:6000) in 5%BSA/0.05% TritonX-100/PBS at 4 °C overnight. Cells were then washed 3X in PBS and incubated with AlexaFluor-594 goat anti-rabbit secondary antibody (Thermo Fisher Scientific; 1:750, Cat #A-11037) in 1%BSA/0.05% TritonX-100/PBS for 45 min at 25 °C. After three washes with PBS, chamber slides were mounted in ProLong Gold with DAPI (Thermo Fisher Scientific). Images were taken using a 63X oil objective and a Zeiss Axio-Imager.Z2 microscope equipped with Zen Blue software (Carl Zeiss Microscopy). Images were obtained as Z-stack sections of 0.2 mm per section containing 18 Z-stacks for each channel. A maximum projection file was generated to identify and count the number of foci and nuclei, and nuclei with >5 RAD51 foci per nucleus were counted as positive. RAD51 foci from 100 nuclei were measured in three to seven independent experiments. Data were processed and plotted using GraphPad Prism 10. Standard deviations were calculated and presented as error bars together with the mean values. *p* values were calculated using Kolmogorov-Smirnov tests.

#### DNA fiber assay

Fiber analysis was used to probe replication tract stability as we have done previously (Kwon et al. 2023; Parplys et al. 2015; Taglialatela et al. 2017). DLD1 cells (EV or expressing full-length BRCA2) were pulse-labeled with 25 μM CldU (20 min), washed three times with PBS, and then pulse-labeled with 250 μM IdU (20 min) in RPMI growth medium. Following IdU labeling, cells were washed with PBS and treated with 2 mM HU (Sigma-Aldrich) or HU + mirin (50⏧μM, Sigma-Aldrich) in growth medium. Labeled cells were collected by scraping, resuspended in ice-cold PBS at 4 × 10^5^ cells/ml and 2 μl of this suspension were spotted onto a glass slide and lysed with 7 μl spreading buffer (0.5% SDS, 200 mM Tris-HCl, pH 7.4, 50 mM EDTA). Slides were incubated for 5 min, then tilted to spread labeled DNA fibers. Next, slides were air-dried and fixed in methanol:acetic acid (3:1), rehydrated in PBS for 10 min and denatured in 2.5 M HCl for 1 h at 21 °C. Slides were then rinsed in PBS and blocked in PBS + 0.1% Triton X-100 (PBS-T) + 5% BSA for 1 h (21 °C). Rat anti-BrdU (1:100, AbD Serotec) and mouse anti-IdU (1:100, Becton Dickinson) were then applied to detect CldU and IdU, respectively. After a 1 h incubation, slides were washed in PBS and stained with Alexa Fluor 488-labeled goat anti-mouse IgG1 antibody and Alexa Fluor 594-labeled goat anti-rat antibody (1:300; Thermo Fisher Scientific). Slides were mounted in Prolong Gold Antifade (Thermo Fisher Scientific) and held at 4 °C until image acquisition. Replication tracts were imaged on a Zeiss Axio-Imager.Z2 microscope equipped with ZEN Blue software (Carl Zeiss Microscopy) using a 63X oil objective. CldU and IdU tracts were measured using ImageJ software. Data are from 3 independent experiments with 150-500 CldU fibers measured. Data were plotted in GraphPad Prism 9. Statistical analysis was conducted using the Kruskal-Wallis test.

## Authors Contributions

F.N. Designed and conducted biochemical assays, designed and purified BRCA2 polypeptides, and wrote the manuscript. Y.K. Provided consistent supervision of biochemical assays and construct design. W.L., W.Z.: Designed and conducted colony survival and CRISPR-Cas9 assays. M.E.U., N.S., C.W.: Designed DLD1 cell lines, designed and carried out fractionation, DNA fiber, and RAD51 foci experiments. S.S.Z.: Designed and carried out RAD51-targeting studies. S.B., R.H., A.M., E.D., D.L., S.O., and E.W. provided key research, materials, and participated in experimental design. Y.K., C.W., C.S., P.S. All contributed to supervising writing and editing of the manuscript.

## Acknowledgements

This study was supported by research grants from the US National Institutes of Health (RO1 CA168635, RO1 ES007061, PO1 CA92584, R35 CA241801 (P.S.); R50CA265315 (Y.K.); R01 GM141091 and R01CA268641 (W.Z.); R01 GM136717, R01 CA23728, R01 CA188347 (A.M.); PO1 CA275717 (A.M., R.H., S.B., P.S., D.L., Y.K., W.Z.), RO1CA205224 (R.H.), R01 GM144579 (C.W.); R01CA246807 (S.B.), Danish Cancer Society (R167-A10921-B224) (C.S.), Cancer Prevention and Research Institute (CPRIT) of Texas awards RP220269 (R.H.) and RP210102 (W.Z.), Congressionally Directed Medical Research Programs award BC191160 (A.M.), ACS Postdoctoral Fellowship PF-22-034-01-DMC (C.M.R.), and NIH predoctoral fellowship awards (F30CA260908, T32CA148724) (F.E.N.). P.S. is the holder of the Robert A. Welch Distinguished Chair in Chemistry (AQ-0012). A.M. is the holder of the Joe R. and Teresa Lozano Long Chair in Cancer and recipient of a Recruitment of Established Investigators Award from CPRIT (RR210023). S.B. is the holder of the Mays Family Foundation Distinguished Chair in Oncology.

## Supplemental Figure Legends

**Figure S1. Alignments highlighting BRCA2 OB-DBD and CTRB-DBD amino acid residues likely involved in DNA binding pertaining to Figure 1.**

**(A)**Comparison of BRCA2 primary sequence across 6 orthologs. We used the NCBI COBALT alignment tool to compare sequences of OB2 **(i)**, OB3 **(ii)**, and the CTRB-DBD **(iii)** BRCA2 ortholog sequences deposited in NCBI. Residues conserved across at least 4 orthologs are indicated by bolded, highlighted single-letter residue abbreviations. **(B)**Purity analysis of wild-type (WT) and mutant variants of BRC4-DBD **(i)** and miniBRCA2 **(ii)**.

**Figure S2. Biochemical analysis for DNA binding pertaining to Figures 2 and 4.**

**(A)**Affinity pulldown to test interaction of RAD51 with BRC4-DBD wild-type and OB fold mutants **(i)**. SDS-PAGE showing BRC4-DBD and RAD51 that remained in the supernatant (S) of the pulldown or in the eluate of resin (E) **(ii). (B)**DNA strand exchange assay to test recombination mediator activity of BRC4-DBD and mutant variants **(i)**. Results were quantified and plotted **(ii)**.

**Figure S3.DNA binding specificity of the OB-DBD and CTRB-DBD pertaining to Figure 3.**

**(A)**EMSA to test binding of ssDNA **(i)** and dsDNA **(ii)** by miniBRCA2 and mutants with the indicated OB fold mutation. **(B)**EMSA to test binding of ssDNA **(i)** and dsDNA **(ii)** by miniBRCA2 and mutants with the OB-DBD^9A^ (DBD-9A), CTRB-DBD^4A^ (CTRB-4A), or OB-DBD^9A^/CTRB-DBD^4A^ (9A+4A) mutation. **(C)**EMSA to test binding of a partial duplex with a 3’ ssDNA overhang by miniBRCA2 and mutants with the OB-DBD^9A^ (DBD-9A), CTRB-DBD^4A^ (CTRB-4A), or OB-DBD^9A^/CTRB-DBD^4A^ (9A+4A) mutation. **(D)**Affinity pulldown was conducted to test miniBRCA2 and mutants with the OB-DBD^9A^ (DBD-9A), CTRB-DBD^4A^ (CTRB-4A), or OB-DBD^9A^/CTRB-DBD^4A^ (9A+4A) mutation for Rad51 interaction. The supernatant (S) and eluate (E) fractions of the pulldown reactions were analyzed by SDS-PAGE with Coomassie blue staining.

**Figure S4.** Clonogenic survival biology and replication fork protection data pertaining to Figures 5 and 6.

**(A)**Western blot of cytoplasmic (C) and nuclear (N) fractions prepared from DLD1 cells expressing wild-type BRCA2 or BRCA2 mutants that harbor the OB-DBD^9A^ (DBD-9A) or CTRB-DBD^4A^ (CTRB-4A) mutation. **(B)**Primary data for the clonogenic survival assays testing DLD1 cell lines of the indicated genotype for sensitivity to MMC, camptothecin, olaparib, and rucaparib. **(C)**Representative immunofluorescence images for RAD51 foci in untreated DLD1 cells of the indicated genotype. **(D)**Representative micrographs of DNA fibers labeled with CldU/IdU in DLD1 cells expressing the indicated BRCA2 species, with the indicated HU (2 mM) and mirin (50 µM) treatments.

## Notes

### Competing Interest Statement

The authors have declared no competing interest.

## References

Belan O, Greenhough L, Kuhlen L, Anand R, Kaczmarczyk A, Gruszka DT, Yardimci H, Zhang X, Rueda DS, West SC, et al. 2023. Visualization of direct and diffusion-assisted RAD51 nucleation by full-length human BRCA2 protein. Mol Cell. 83(16):2925-2940.e8. doi:10.1016/j.molcel.2023.06.031.

Bhat KP, Cortez D. 2018. RPA and RAD51: Fork reversal, fork protection, and genome stability. Nat Struct Mol Biol. 25(6):446–453. doi:10.1038/s41594-018-0075-z. 10.1038/s41594-018-0075-z.

Bochkarev A, Bochkareva E, Frappier L, Edwards AM. 1999. The crystal structure of the complex of replication protein A subunits RPA32 and RPA14 reveals a mechanism for single-stranded DNA binding. EMBO Journal. 18(16):4498–4504. doi:10.1093/emboj/18.16.4498.

Carreira A, Hilario J, Amitani I, Baskin RJ, Shivji MKK, Venkitaraman AR, Kowalczykowski SC. 2009. The BRC Repeats of BRCA2 Modulate the DNA-Binding Selectivity of RAD51. Cell. 136(6):1032–1043. doi:10.1016/j.cell.2009.02.019. 10.1016/j.cell.2009.02.019.

Cejka P, Symington LS. 2021. DNA End Resection: Mechanism and Control. Annu Rev Genet. 55:285–307. doi:10.1146/annurev-genet-071719-020312.

Chalermrujinanant C, Michowski W, Sittithumcharee G, Esashi F, Jirawatnotai S. 2016. Cyclin D1 promotes BRCA2-Rad51 interaction by restricting cyclin A/B-dependent BRCA2 phosphorylation. Oncogene. 35(22):2815–2823. doi:10.1038/onc.2015.354.

Chatterjee G, Jimenez-Sainz J, Presti T, Nguyen T, Jensen RB. 2016. Distinct binding of BRCA2 BRC repeats to RAD51 generates differential DNA damage sensitivity. Nucleic Acids Res. 44(11):5256–5270. doi:10.1093/nar/gkw242.

Cortez D. 2019. Replication-Coupled DNA Repair. Mol Cell. 74(5):866–876. doi:10.1016/j.molcel.2019.04.027.

Daley JM, Niu H, Miller AS, Sung P. 2015. Biochemical mechanism of DSB end resection and its regulation. DNA Repair (Amst). 32:66–74. doi:10.1016/j.dnarep.2015.04.015.

Davies OR, Pellegrini L. 2007. Interaction with the BRCA2 C terminus protects RAD51-DNA filaments from disassembly by BRC repeats. Nat Struct Mol Biol. 14(6):475–483. doi:10.1038/nsmb1251.

Donoho G, Brenneman MA, Cui TX, Donoviel D, Vogel H, Goodwin EH, Chen DJ, Hasty P. 2003. Deletion of Brca2 exon 27 causes hypersensitivity to DNA crosslinks, chromosomal instability, and reduced life span in mice. Genes Chromosomes Cancer. 36(4):317–331. doi:10.1002/gcc.10148.

Esashi F, Christ N, Cannon J, Liu Y, Hunt T, Jasin M, West SC. 2005. CDK-dependent phosphorylation of BRCA2 as a regulatory mechanism for recombinational repair. Nature. 434(7033):598–604. doi:10.1038/nature03404.

Esashi F, Galkin VE, Yu X, Egelman EH, West SC. 2007. Stabilization of RAD51 nucleoprotein filaments by the C-terminal region of BRCA2. Nat Struct Mol Biol. 14(6):468–474. doi:10.1038/nsmb1245.

Fan J, Pavletich NP. 2012. Structure and conformational change of a replication protein A heterotrimer bound to ssDNA. Genes Dev. 26(20):2337–2347. doi:10.1101/gad.194787.112.

Filippo JS, Chi P, Sehorn MG, Etchin J, Krejci L, Sung P. 2006. Recombination mediator and Rad51 targeting activities of a human BRCA2 polypeptide. Journal of Biological Chemistry. 281(17):11649–11657. doi:10.1074/jbc.M601249200.

Gradia SD, Ishida JP, Tsai MS, Jeans C, Tainer JA, Fuss JO. 2017. MacroBac: New Technologies for Robust and Efficient Large-Scale Production of Recombinant Multiprotein Complexes. In: Methods in Enzymology. Vol. 592. Academic Press Inc. p. 1–26.

Halder S, Sanchez A, Ranjha L, Acharya A, Halder S, Sanchez A, Ranjha L, Reginato G, Ceppi I, Acharya A. 2022. Article Double-stranded DNA binding function of RAD51 in DNA protection and its regulation by BRCA2 Article Double-stranded DNA binding function of RAD51 in DNA protection and its regulation by BRCA2. Mol Cell. 82:1–13. doi:10.1016/j.molcel.2022.08.014. 10.1016/j.molcel.2022.08.014.

Henricksen LA, Umbricht CB, Wold MS. 1994. Recombinant replication protein A: Expression, complex formation, and functional characterization. Journal of Biological Chemistry. 269(15):11121–11132. doi:10.1016/s0021-9258(19)78100-9.

Huang, Y., Li, W., Foo, T., Ji, J.-H., Wu, B., Tomimatsu, N., Fang, Q., Gao, B., Long, M., Xu, J., Maqbool, R., Mukherjee, B., Ni, T., Alejo, S., He, Y., Burma, S., Lan, L., Xia, B., & Zhao, W. (2024). DSS1 restrains BRCA2’s engagement with dsDNA for homologous recombination, replication fork protection, and R-loop homeostasis. Nature Communications, 15(1), 7081. 10.1038/s41467-024-51557-6

Jasin M. 2002. Homologous repair of DNA damage and tumorigenesis: The BRCA connection. Oncogene. 21(58 REV. ISS. 8):8981–8993. doi:10.1038/sj.onc.1206176.

Jensen RB, Carreira A, Kowalczykowski SC. 2010. Purified human BRCA2 stimulates RAD51-mediated recombination. Nature. 467(7316):678–683. doi:10.1038/nature09399.

Jimenez-Sainz, J., Mathew, J., Moore, G., Lahiri, S., Garbarino, J., Eder, J. P., Rothenberg, E., & Jensen, R. B. (2022). BRCA2 BRC missense variants disrupt RAD51-dependent DNA repair. ELife, 11. 10.7554/eLife.79183

Kaminski N, Wondisford AR, Kwon Y, Lynskey ML, Bhargava R, Barroso-González J, García-Expósito L, He B, Xu M, Mellacheruvu D, et al. 2022. RAD51AP1 regulates ALT-HDR through chromatin-directed homeostasis of TERRA. Mol Cell. 82(21):4001-4017.e7. doi:10.1016/j.molcel.2022.09.025.

Kass EM, Lim PX, Helgadottir HR, Moynahan ME, Jasin M. 2016. ARTICLE Robust homology-directed repair within mouse mammary tissue is not specifically affected by Brca2 mutation. doi:10.1038/ncomms13241.

Kim TM, Son MY, Dodds S, Hu L, Hasty P. 2014. Deletion of BRCA2 exon 27 causes defects in response to both stalled and collapsed replication forks. Mutation Research - Fundamental and Molecular Mechanisms of Mutagenesis. 766–767:66–72. doi:10.1016/j.mrfmmm.2014.06.003. 10.1016/j.mrfmmm.2014.06.003.

Konstantinopoulos PA, Ceccaldi R, Shapiro GI, D’Andrea AD. 2015. Homologous recombination deficiency: Exploiting the fundamental vulnerability of ovarian cancer. Cancer Discov. 5(11):1137–1154. doi:10.1158/2159-8290.CD-15-0714.

Kwon Y, Rösner H, Zhao W, Selemenakis P, He Z, Kawale AS, Katz JN, Rogers CM, Neal FE, Shabestari AB, et al. 2023. DNA binding and RAD51 engagement by the BRCA2 C-terminus orchestrate DNA repair and replication fork preservation. Nat Commun. 14(432). doi:10.1038/s41467-023-36211-x.

Kwon Y, Zhao W, Sung P. 2011. Biochemical Studies on Human Rad51-Mediated Homologous Recombination. Methods Mol Biol. 745:421–435. doi:10.1007/978-1-61779-129-1.

Lo T, Pellegrini L, Venkitaraman AR, Blundell TL. 2003. Sequence fingerprints in BRCA2 and RAD51: Implications for DNA repair and cancer. DNA Repair (Amst). 2(9):1015–1028. doi:10.1016/S1568-7864(03)00097-1.

McAllister KA, Bennett LM, Houle CD, Ward T, Malphurs J, Collins NK, Cachafeiro C, Haseman J, Goulding EH, Bunch D, et al. 2002. Cancer susceptibility of mice with a homozygous deletion in the COOH-terminal domain of the Brca2 gene. Cancer Res. 62(4):990–994.

Mijic S, Zellweger R, Chappidi N, Berti M, Jacobs K, Mutreja K, Ursich S, Ray Chaudhuri A, Nussenzweig A, Janscak P, et al. 2017. Replication fork reversal triggers fork degradation in BRCA2-defective cells. Nat Commun. 8(1):1–11. doi:10.1038/s41467-017-01164-5.

Morimatsu M, Donoho G, Hasty P. 1998. Cells deleted for Brca2 COOH terminus exhibit hypersensitivity to γ-radiation and premature senescence. Cancer Res. 58(15):3441–3447.

Moynahan ME, Pierce AJ, Jasin M. 2001. BRCA2 is required for homology-directed repair of chromosomal breaks. Mol Cell. 7(2):263–272. doi:10.1016/S1097-2765(01)00174-5.

Parplys AC, Zhao W, Sharma N, Groesser T, Liang F, Maranon DG, Leung SG, Grundt K, Idate R, Østvold AC, et al. 2015. NUCKS1 is a novel RAD51AP1 paralog important for homologous recombination and genome stability. 43(20):9817–9834. doi:10.1093/nar/gkv859.

Pinder J, Salsman J, Dellaire G. 2015. Nuclear domain “knock-in” screen for the evaluation and identification of small molecule enhancers of CRISPR-based genome editing. Nucleic Acids Res. 43(19):9379–9392. doi:10.1093/nar/gkv993.

Prakash R, Zhang Y, Feng W, Jasin M. 2015. Homologous recombination and human health: The roles of BRCA1, BRCA2, and associated proteins. Cold Spring Harb Perspect Biol. 7(4):1–28. doi:10.1101/cshperspect.a016600.

Rajagopalan S, Andreeva A, Rutherford TJ, Fersht AR. 2010. Mapping the physical and functional interactions between the tumor suppressors p53 and BRCA2. Proc Natl Acad Sci U S A. 107(19):8587–8592. doi:10.1073/pnas.1003689107.

San Filippo J, Sung P, Klein H. 2008. Mechanism of eukaryotic homologous recombination. Annu Rev Biochem. 77:229–257. doi:10.1146/annurev.biochem.77.061306.125255.

Schlacher K, Christ N, Siaud N, Egashira A, Wu H, Jasin M. 2011. Double-strand break repair-independent role for BRCA2 in blocking stalled replication fork degradation by MRE11. Cell. 145(4):529–542. doi:10.1016/j.cell.2011.03.041.

Scully R, Panday A, Elango R, Willis NA. 2019. DNA double-strand break repair-pathway choice in somatic mammalian cells. Nat Rev Mol Cell Biol. 20(11):698–714. doi:10.1038/s41580-019-0152-0.

Sharan SK, Morimatsu M, Albrecht U, Lim DS, Regel E, Dinh C, Sands A, Eichele G, Hasty P, Bradley A. 1997. Embryonic lethality and radiation hypersensitivity mediated by Rad51 in mice lacking Brca2. Nature. 386(6627):804–810.

Shivji MKK, Mukund SR, Rajendra E, Chen S, Short JM, Savill J, Klenerman D, Venkitaraman AR. 2009. The BRC repeats of human BRCA2 differentially regulate RAD51 binding on single-versus double-stranded DNA to stimulate strand exchange. Proc Natl Acad Sci U S A. 106(32):13254–13259. doi:10.1073/pnas.0906208106.

Siaud N, Barbera MA, Egashira A, Lam I, Christ N, Schlacher K, Xia B, Jasin M. 2011. Plasticity of BRCA2 function in homologous recombination: Genetic interactions of the PALB2 and DNA binding domains. PLoS Genet. 7(12). doi:10.1371/journal.pgen.1002409.

Sigurdsson S, Trujillo K, Song BW, Stratton S, Sung P. 2001. Basis for Avid Homologous DNA Strand Exchange by Human Rad51 and RPA. Journal of Biological Chemistry. 276(12):8798–8806. doi:10.1074/jbc.M010011200.

Suzuki K, Bose P, Leong-Quong RY, Fujita DJ, Riabowol K. 2010. REAP: A two minute cell fractionation method. BMC Res Notes. 3. doi:10.1186/1756-0500-3-294.

Taglialatela A, Alvarez S, Leuzzi G, Sannino V, Ranjha L, Huang JW, Madubata C, Anand R, Levy B, Rabadan R, et al. 2017. Restoration of Replication Fork Stability in BRCA1- and BRCA2-Deficient Cells by Inactivation of SNF2-Family Fork Remodelers. Mol Cell. 68(2):414-430.e8. doi:10.1016/j.molcel.2017.09.036. 10.1016/j.molcel.2017.09.036.

Theobald DL, Mitton-Fry RM, Wuttke DS. 2003. Nucleic acid recognition by OB-fold proteins. Annu Rev Biophys Biomol Struct. 32:115–133. doi:10.1146/annurev.biophys.32.110601.142506.

Tye S, Ronson GE, Morris JR. 2021. A fork in the road: Where homologous recombination and stalled replication fork protection part ways. Semin Cell Dev Biol. 113(May 2020):14–26. doi:10.1016/j.semcdb.2020.07.004.

Yang H, Jeffrey PD, Miller J, Kinnucan E, Sun Y, Thomä NH, Zheng N, Chen PL, Lee WH, Pavletich NP. 2002. BRCA2 function in DNA binding and recombination from a BRCA2-DSS1-ssDNA structure. Science (1979). 297(5588):1837–1848. doi:10.1126/science.297.5588.1837.

Zhao W, Vaithiyalingam S, San Filippo J, Maranon DG, Jimenez-Sainz J, Fontenay G V., Kwon Y, Leung SG, Lu L, Jensen RB, et al. 2015. Promotion of BRCA2-Dependent Homologous Recombination by DSS1 via RPA Targeting and DNA Mimicry. Mol Cell. 59(2):176–187. doi:10.1016/j.molcel.2015.05.032. 10.1016/j.molcel.2015.05.032.

Zhao W, Wiese C, Kwon Y, Hromas R, Sung P. 2019. The BRCA tumor suppressor network in chromosome damage repair by homologous recombination. Annu Rev Biochem. 88:221–245. doi:10.1146/annurev-biochem-013118-111058.

