## Supplemental Figures and Tables for "Distinct roles of the two BRCA2 DNA binding domains in DNA damage repair and replication fork preservation"

### Slide 1
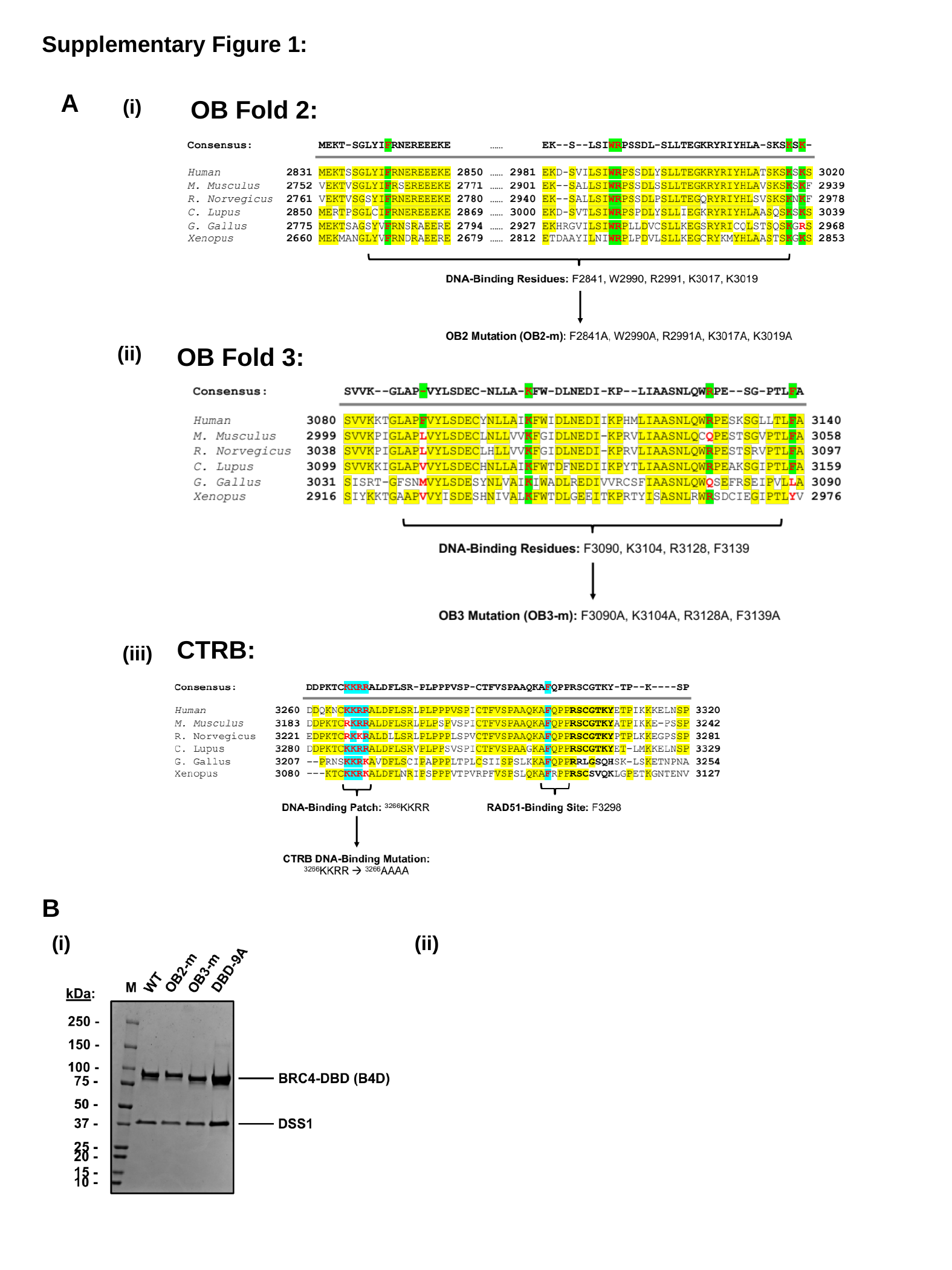

Supplementary Figure 1:
A
OB Fold 2:
(i)
(ii)
OB Fold 3:
CTRB:
(iii)
B
(i)
(ii)

### Slide 2
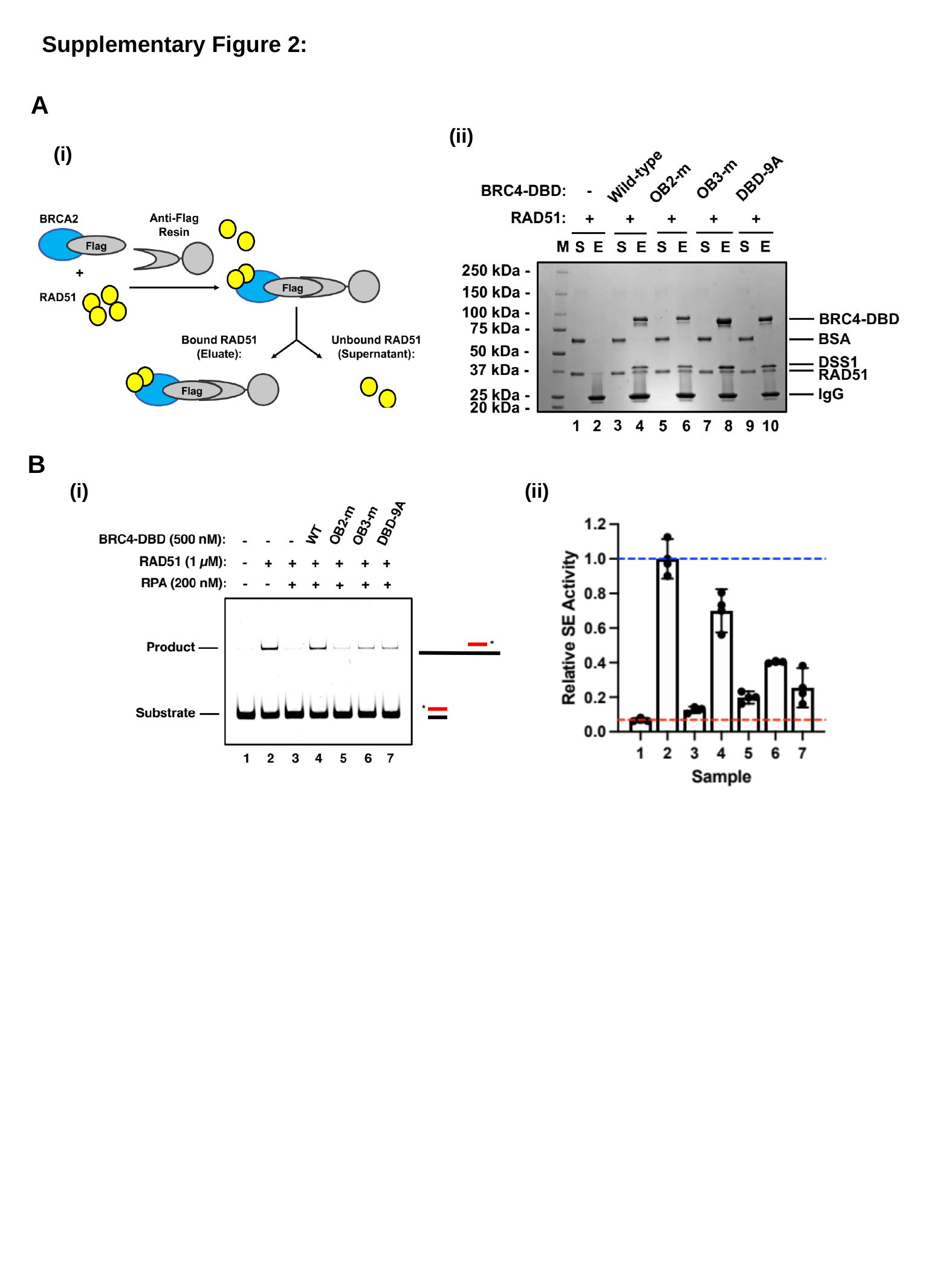

Supplementary Figure 2:
A
(ii)
(i)
B
(i)
(ii)

### Slide 3
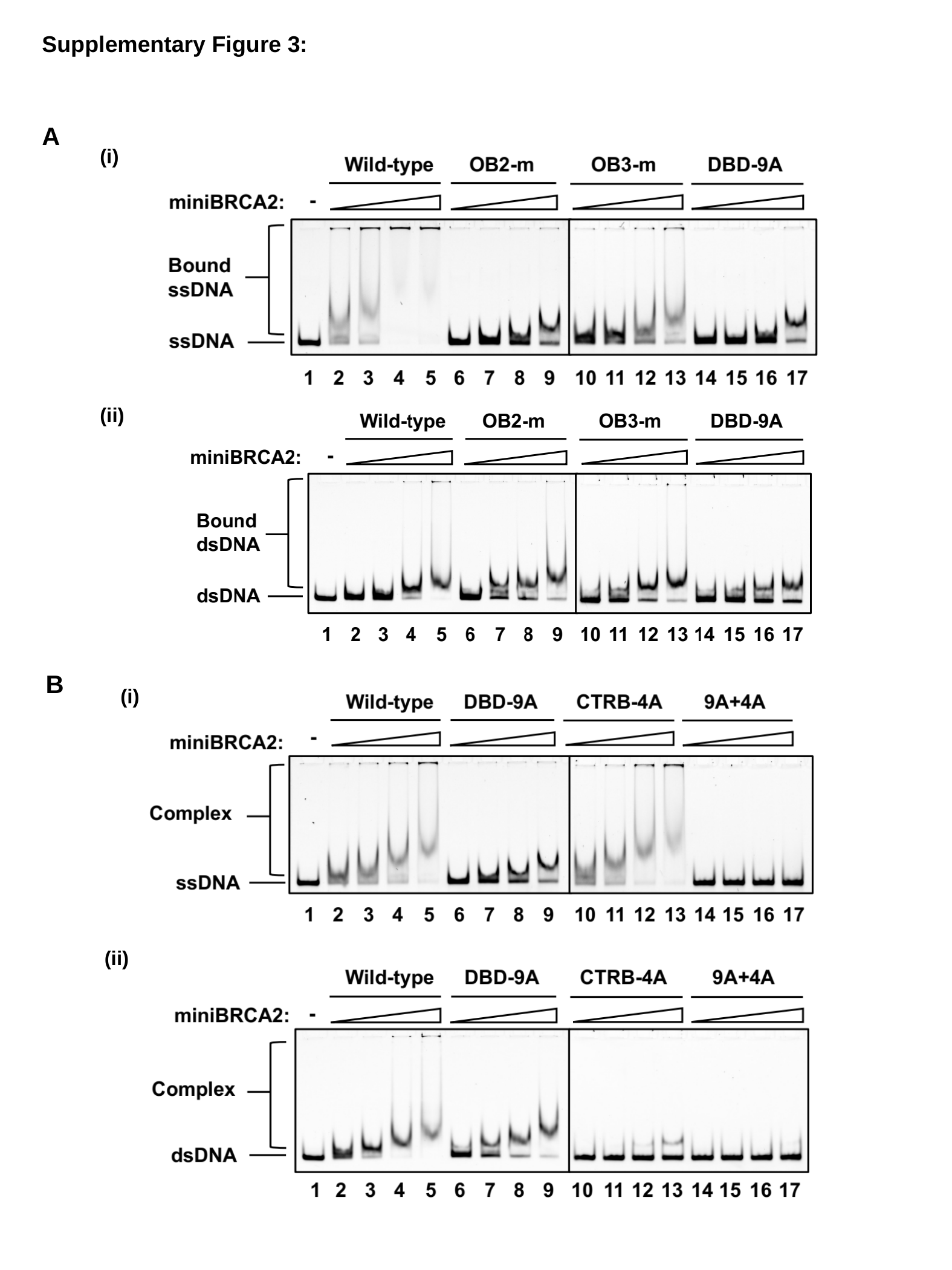

Supplementary Figure 3:
A
(i)
(ii)
B
(i)
(ii)

### Slide 4
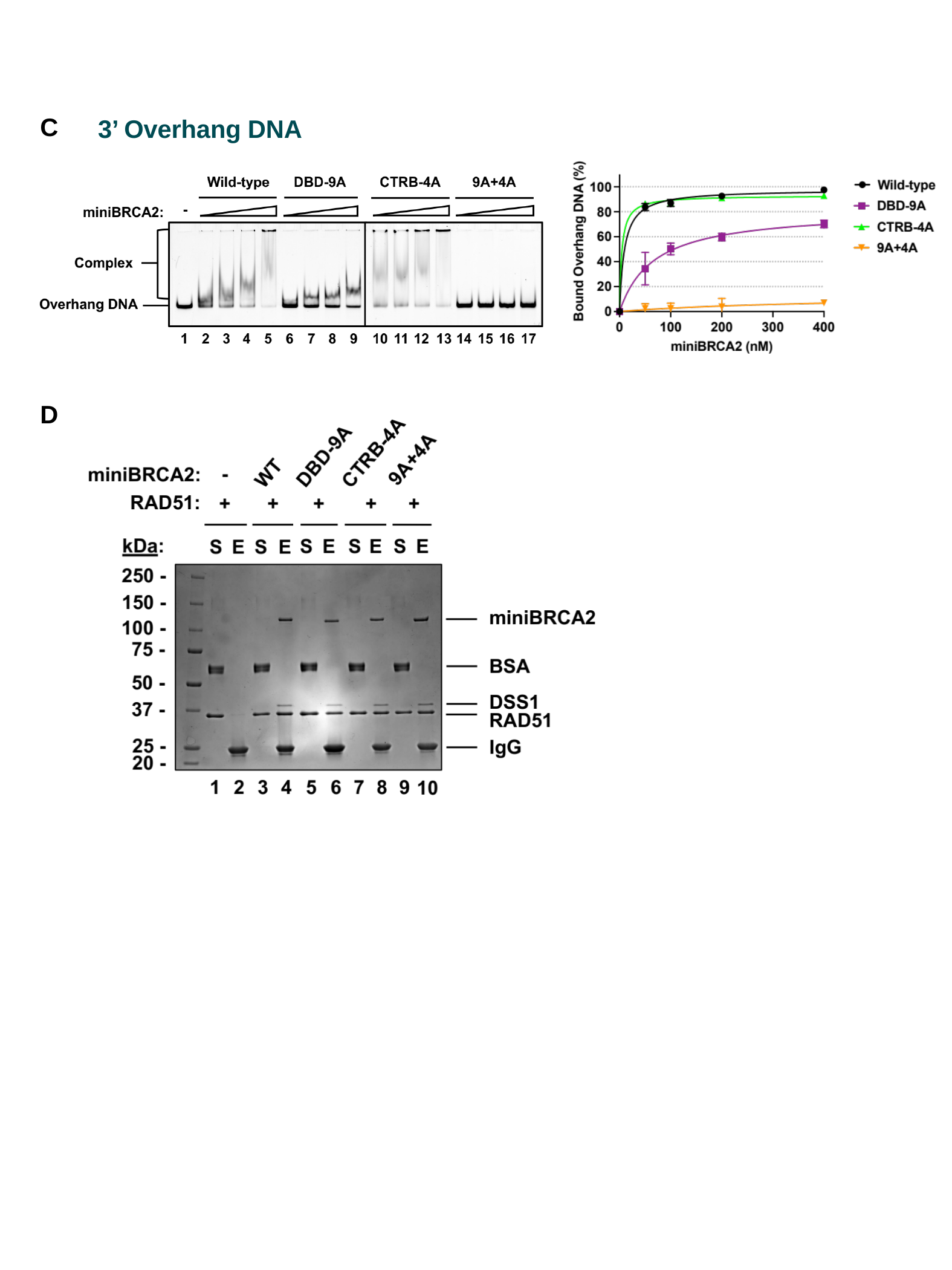

C
3’ Overhang DNA
D

### Slide 5
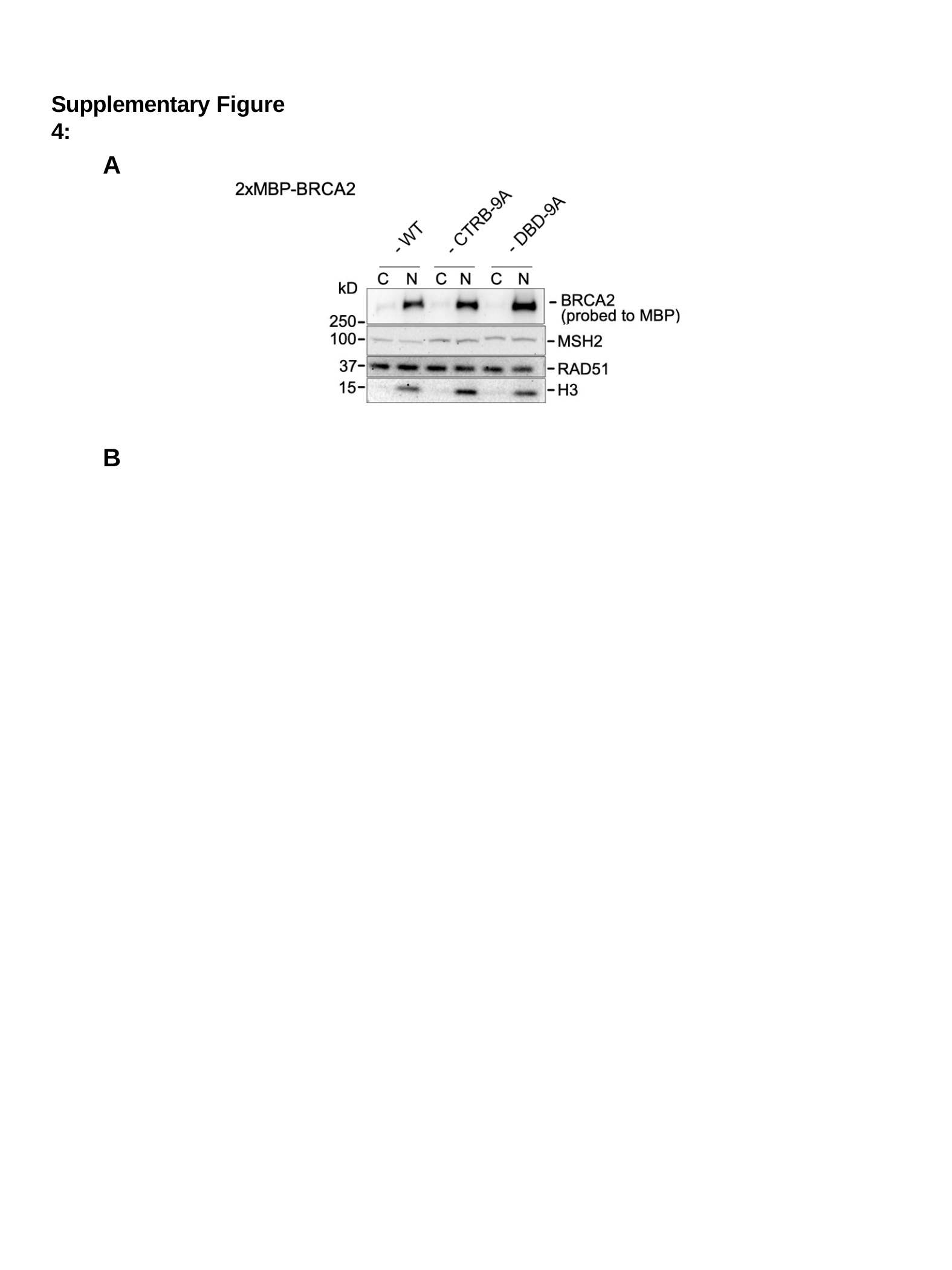

Supplementary Figure 4:
A
B

### Slide 6
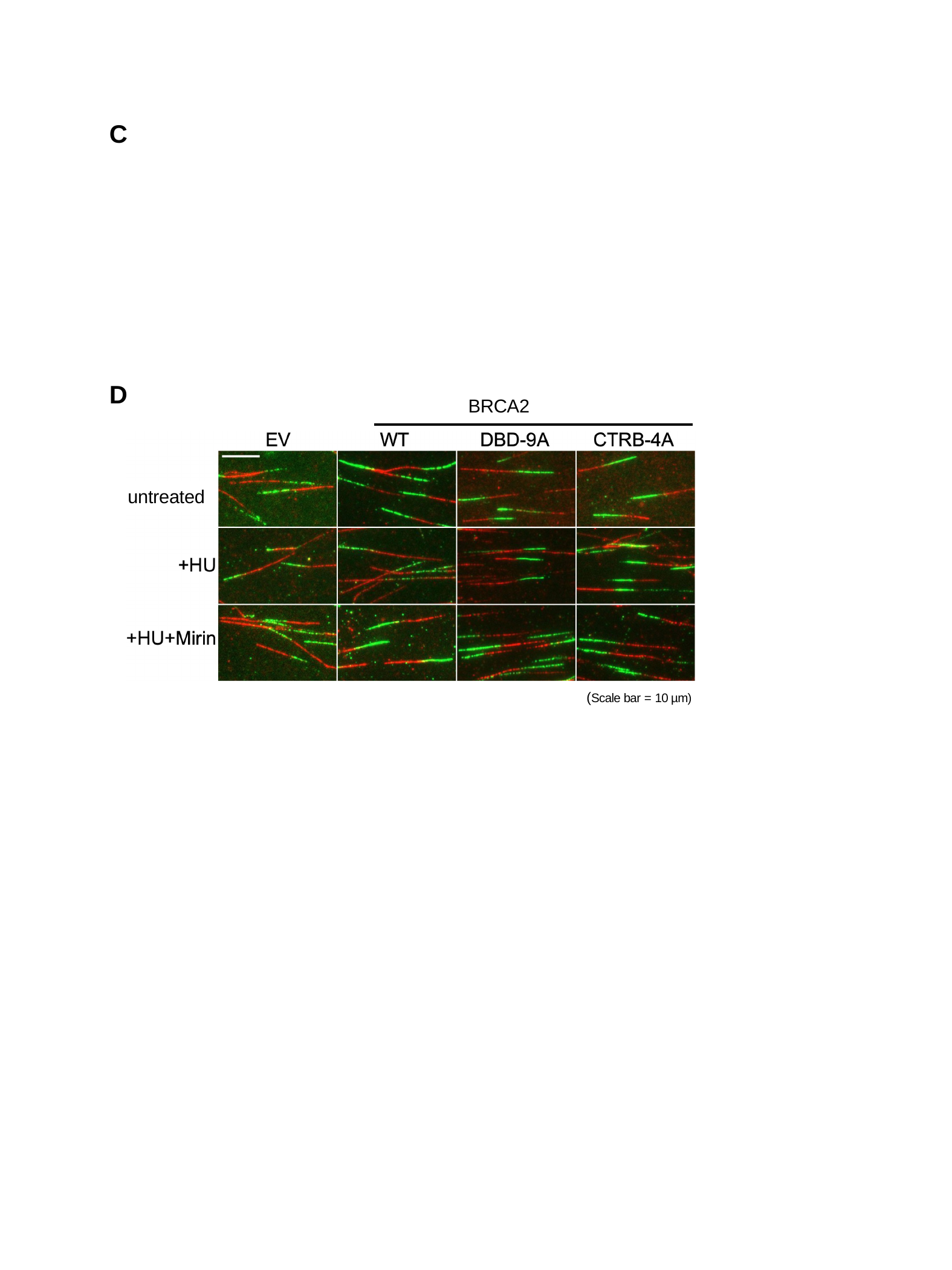

C
D
BRCA2
untreated
(Scale bar = 10 µm)

### Slide 7
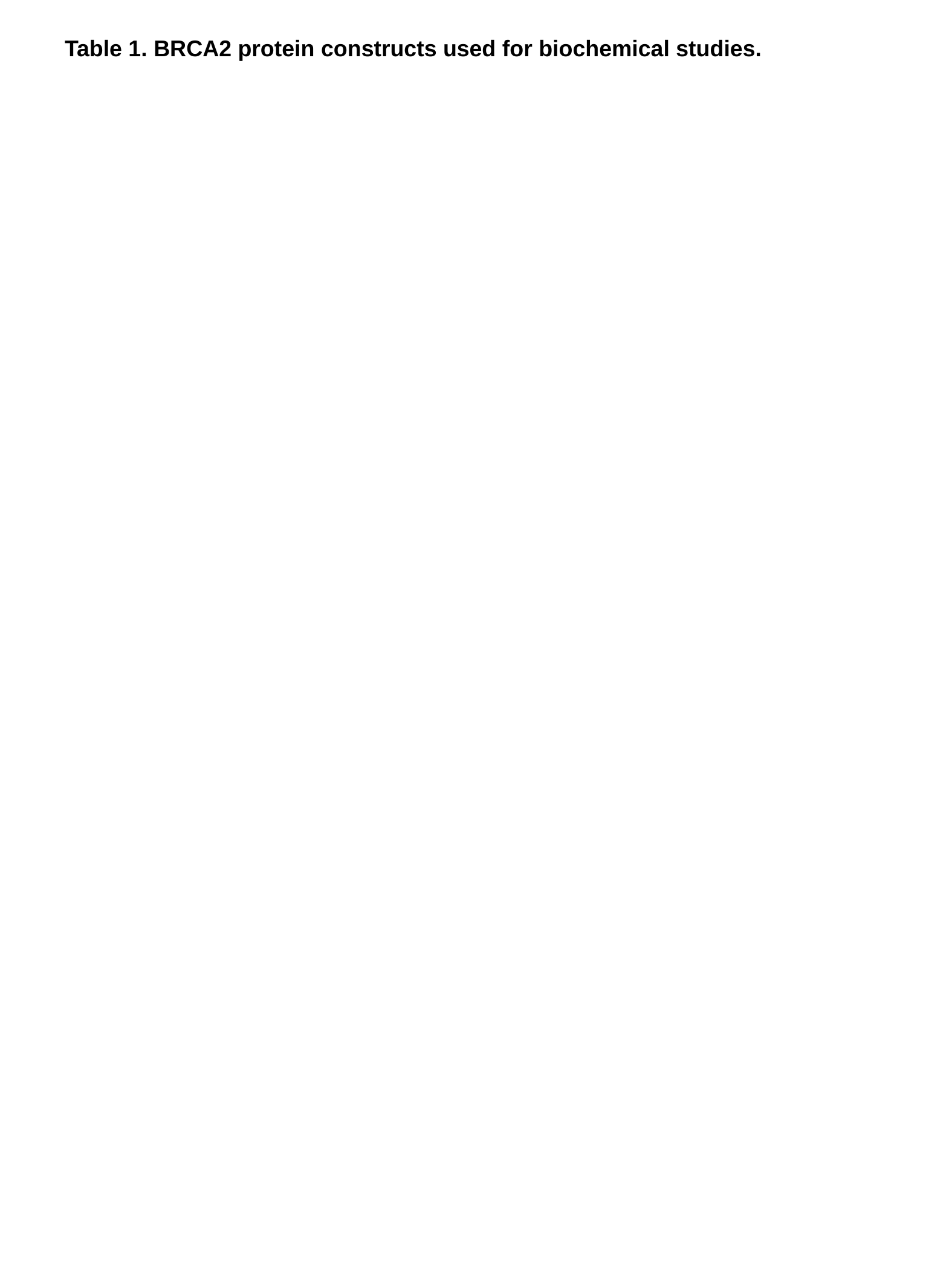

#### Table 1. BRCA2 protein constructs used for biochemical studies.

### Slide 8
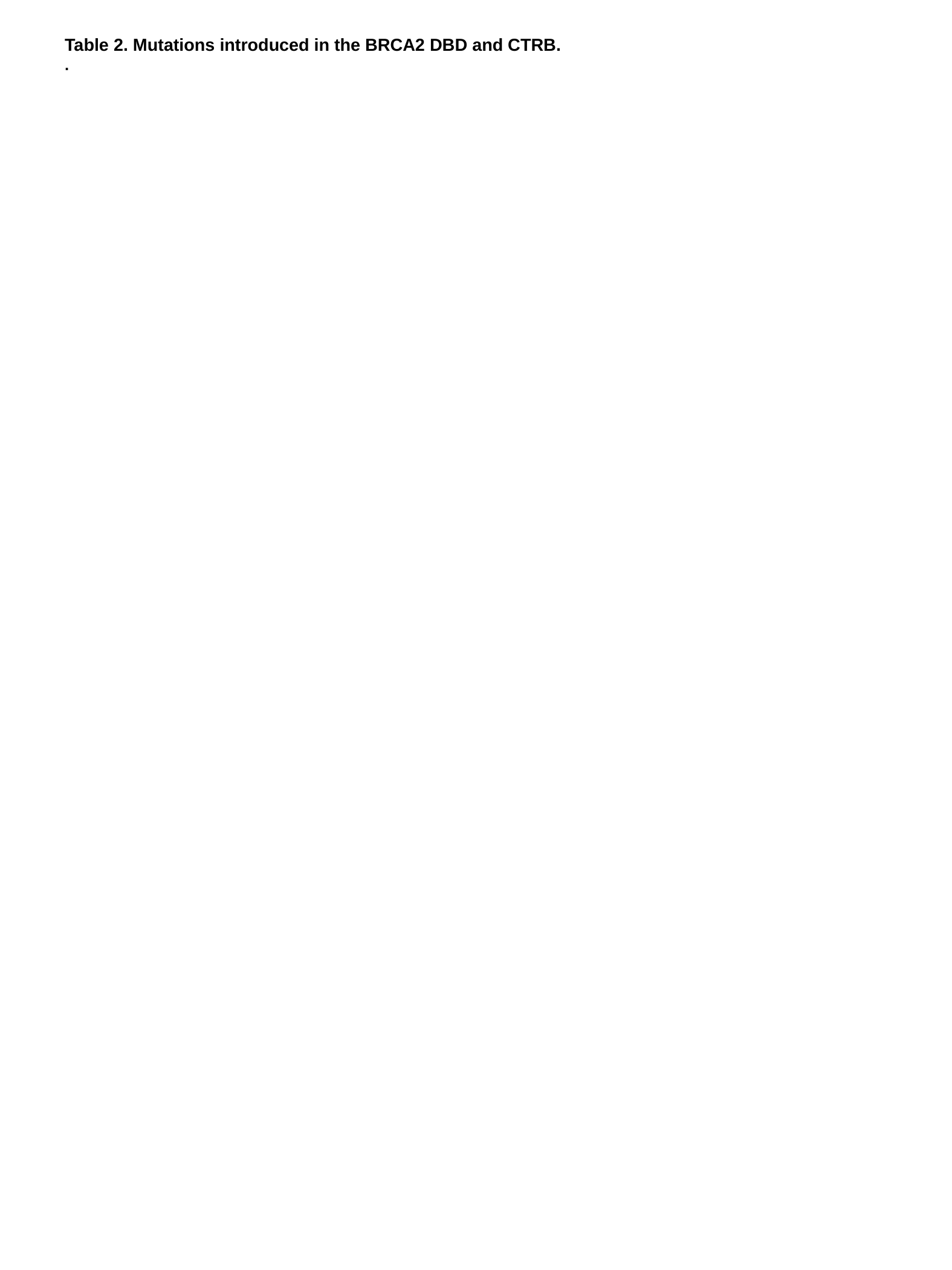

#### Table 2. Mutations introduced in the BRCA2 DBD and CTRB. .

### Slide 9
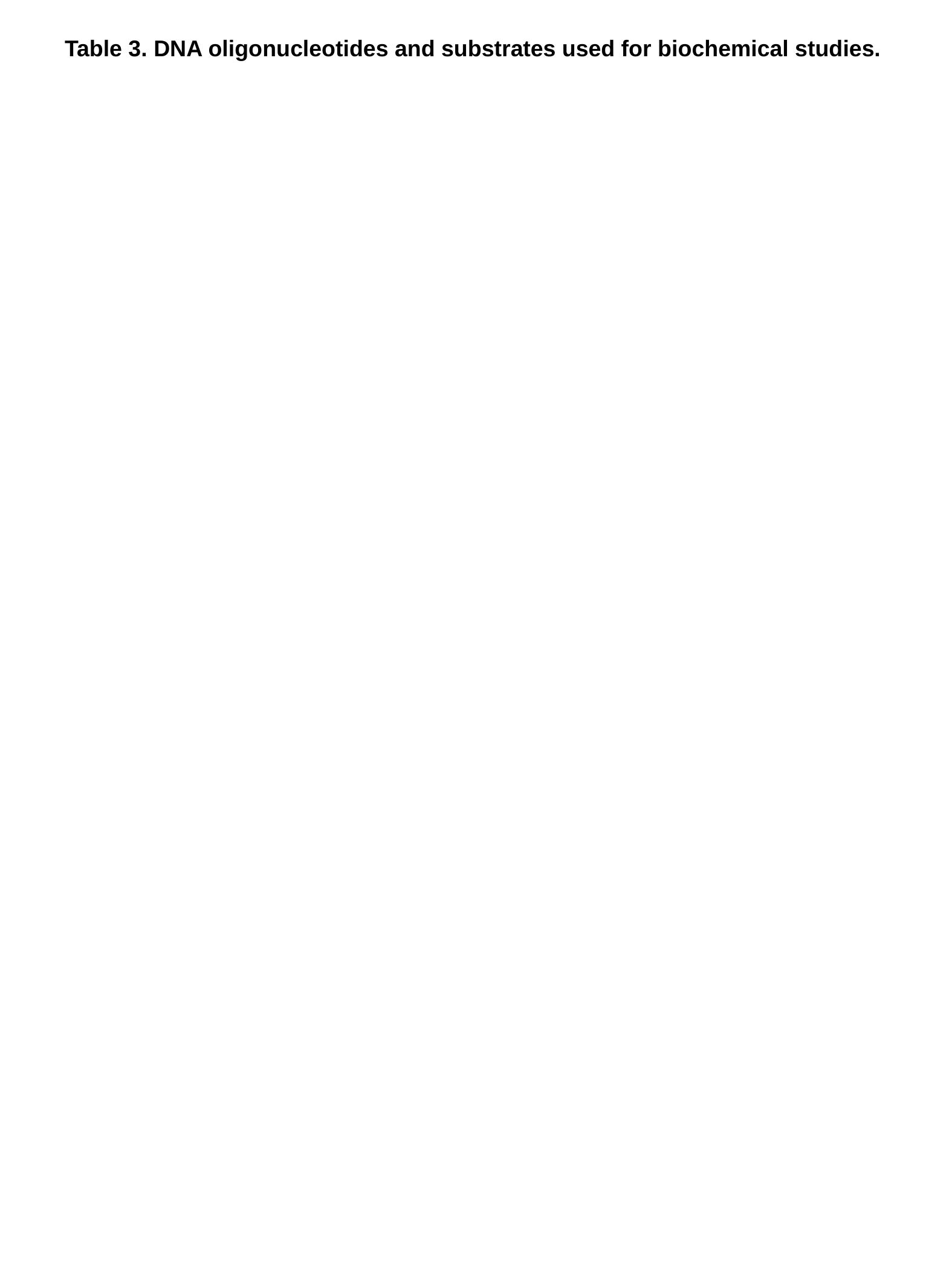

#### Table 3. DNA oligonucleotides and substrates used for biochemical studies.
